# Characterization of the Endometrial Transcriptome in Early Diestrus Influencing Pregnancy Establishment in Dairy Cattle after Transfer of In-Vitro Produced Embryos

**DOI:** 10.1101/2020.03.04.977082

**Authors:** Gianluca Mazzoni, Hanne S Pedersen, Maria B Rabaglino, Poul Hyttel, Henrik Callesen, Haja N Kadarmideen

**Affiliations:** Department of Veterinary and Animal Sciences, Faculty of Health and Medical Sciences, University of Copenhagen, Grønnegårdsvej 7, 1870 Frederiksberg C, Denmark; Department of Health Technology, Technical University of Denmark, Ørsteds Plads, Building 345C, DK-2800 Kgs. Lyngby, Denmark; Department of Animal Science, Aarhus University, Blichers Alle 20, 8830 Tjele, Denmark; Quantitative Genetics, Bioinformatics and Computational Biology Group, Department of Applied Mathematics and Computer Science, Technical University of Denmark, Kemitorvet, 2800 Kgs. Lyngby, Denmark

**Keywords:** RNASeq, Fertility, Candidate genes, Biomarkers, Molecular Pathways

## Abstract

Modifications of the endometrial transcriptome at day 7 of the estrus cycle are crucial to maintain gestation after transfer of in-vitro produced (IVP) embryos. The aim of this study was to identify genes, and their related biological mechanisms, in the endometria of recipient lactating dairy cows that become pregnant in the subsequent estrus cycle upon transfer of IVP embryos. Endometrial biopsies were taken from lactating Holstein Friesian cows on day 6-8 of the estrus cycle followed by embryo transfer in the following cycle. Animals were classified retrospectively as pregnant (PR, n=8) or non-pregnant (non-PR, n=11) cows, according to pregnancy status at 26-47 days. Extracted mRNAs from endometrial samples were sequenced with an Illumina platform to determine differentially expressed genes (DEG) between the endometrial transcriptome from PR and non-PR cows. There were 111 DEG (FDR<0.05), which were mainly related to extracellular matrix interaction, histotroph metabolic composition, prostaglandin synthesis, TGF-β signaling as well as inflammation and leukocyte activation. Comparison of these DEG with DEG identified in two public external datasets confirmed the more fertile endometrial molecular profile of PR cows. In conclusion, this study provides insights into the key early endometrial mechanisms for pregnancy establishment, after IVP embryo transfer in dairy cows.

## Introduction

The fertilization rate after artificial insemination (AI) of dairy cattle is estimated to be 80-90% whereas the average calving rate is 55% ^1,2^. This high pregnancy loss occurs during the first part of the embryonic period ^3^, mainly between days 7 and 16 of gestation ^4^, due mostly to the inability of the uterus to support embryo implantation and growth ^5^.

Successful establishment and maintenance of a pregnancy is dependent on both embryonic viability and the uterine environment sustaining embryonic development. With respect to embryonic viability, transfer of *in-vitro* produced (IVP) embryos generally results in lower pregnancy rates (30-40%) as compared with embryos produced *in-vivo* after superovulation and AI of the donor cow (55–60%) ^6,7^. IVP embryo quality is dependent on oocyte competence, sperm quality, and in-vitro handling, and despite many efforts to improve media and methodologies, IVP and *in-vivo* produced embryos still deviate in several ways ^8–10^, due to which optimization of the IVP process is subject to intense investigation yet (reviewed by Lonergan ^11^). With respect to uterine environment, estradiol produced by follicles around estrus primes the endometrium ^12^, and progesterone produced by the corpus luteum (CL) in diestrus induces endometrial receptivity ^13^, which is known as the period when the uterus environment is physiologically ideal for establishment of pregnancy. During diestrus, the endometrial stroma is thickened and uterine glands enlarged, resulting in an increased production of histotroph ^14,15^ that is absorbed by the trophoblast of the potential embryo ^16^. The thickness of the endometrium depends on the concentration of circulating estradiol and progesterone, and it has been shown to vary between natural or induced estrous ^17^.

Endometrial changes occur in every reproductive cycle, independent of the presence of a conceptus ^18^. Interestingly, the endometrial transcriptome, underlying these changes, may convey a higher endometrial receptivity in some animals than in others. Hence, the endometrial transcriptome at day 7 differs between heifers that maintain the pregnancy and those that do not ^19^. On top of these cyclic changes in endometrial transcriptome, signals from an embryo further modify endometrial gene expression already from day 7 in the estrus cycle ^20^. Taken together, the endometrial transcriptome varies over the estrus cycle, between receptive and less receptive individuals, and with respect to whether an embryo is present or not.

Regarding endometrial receptivity, experiments performed in Nellore cows and Simmental heifers ^19,21,22^, after estrus synchronization, have unveiled important differences in the endometrial transcriptome at day 6 or 7 of animals that become pregnant after AI or embryo transfer (ET), respectively, compared to the non-pregnant ones. However, disparities in embryo survival between heifers and lactating cows ^2^, the different effects of progesterone on the fertility of dairy and beef cows ^23^ and the variations in hormonal concentration and uterine thickness between induced or natural estrus ^17^, indicate the need of further studies for a better comprehension of the mechanisms controlling endometrial receptivity.

The aim of this study was to determine biological processes and pathways that convey receptivity in the endometrium for establishment of pregnancy based on the identification of differentially expressed genes (DEG) in the endometrium on day 6-8 in diestrus of recipient lactating dairy cows, that become pregnant in the subsequent estrus cycle after transfer of IVP embryos, compared to those that do not establish pregnancy. Thus, cows were classified retrospectively as pregnant (PR) or non-pregnant (non-PR) according to their status.

Furthermore, results obtained from the present study were compared to the endometrial transcriptome differentially expressed at day 7 between cows with high or low merit for fertility ^24^ or cows with high or lower levels of progesterone ^25^. The goal was to employ a bioinformatic approach to test the hypothesis that the observed differences in the transcriptome are attributed more to the inherent endometrial molecular signature of the animal than to hormonal-and other-variations between cycles.

## Results

### RNA-Seq data pre-processing and alignment

mRNA samples extracted from the endometrial biopsies were paired-end sequenced with the Illumina HiSeq 2500 platform, except for one sample from the non-PR group that was excluded due to low RNA quality. The sequencing generated, on average, 44,252,601 read pairs per sample and 41,799,192 of the read pairs (94% of the read pairs) were retained after the pre-processing step. Among them, on average, 91% were uniquely mapped to the bovine reference genome (43% of the read pairs mapped to exonic regions, 27% of the read pairs mapped to intronic regions and 30% of the read pairs mapped to intergenic regions; Supplementary Table 1)

### Expression profile clustering and differential expression analysis

We identified 111 DEG (FDR < 0.05) in PR vs. non-PR cows. Among them, 60 were up-and 51 were down-regulated DEG in PR cows compared to non-PR cows (Supplementary Table 1). The hierarchical clustering based on the expression profiles (Fig. 1) showed a clear association of the samples from PR and non-PR cows, although two samples from non-PR cows clustered with the samples from the PR group.

**Figure 1.**
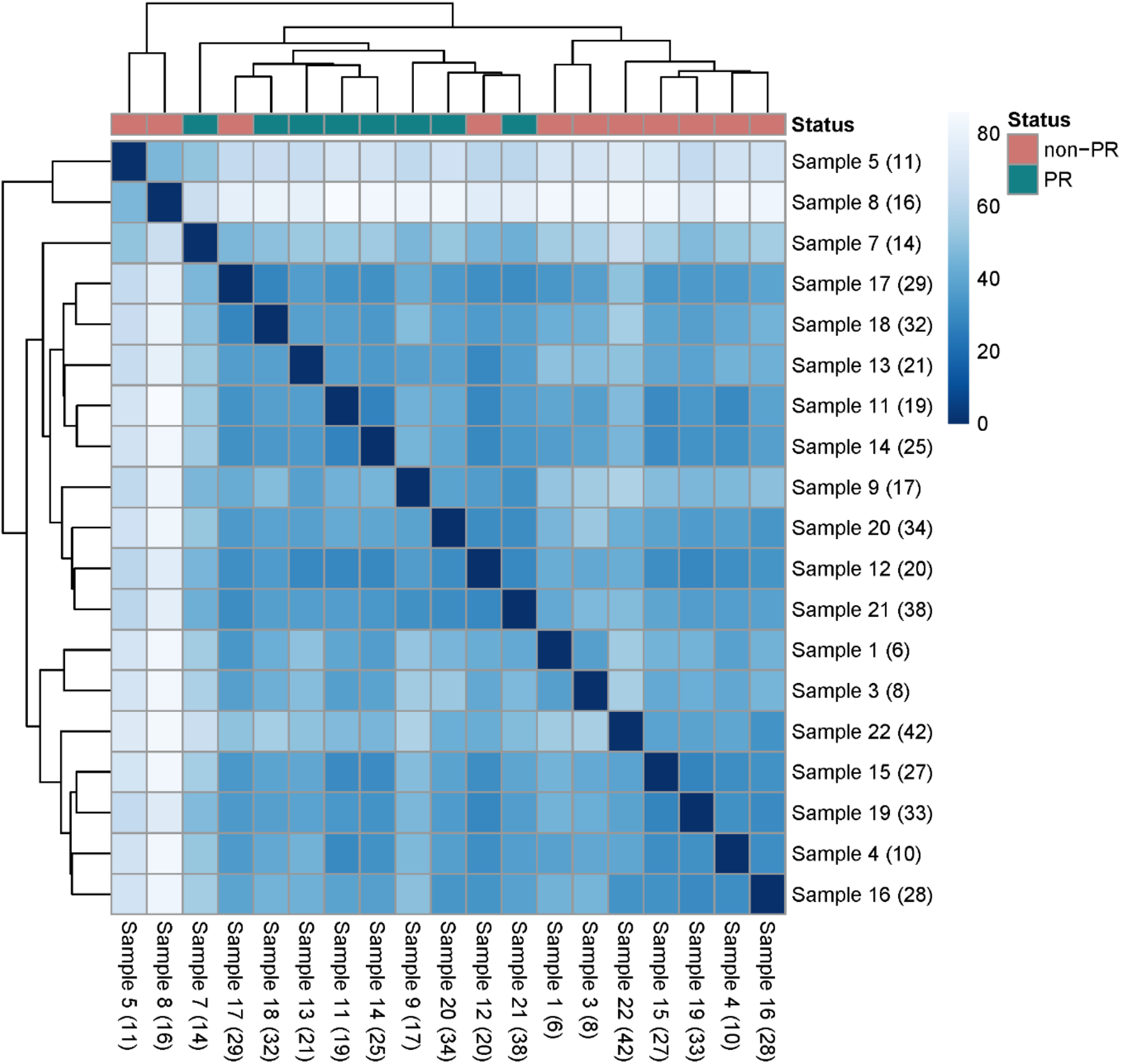
Hierarchical clustering based on the distances of the correlation between expression profiles across samples. The pregnant (PR) cows (red dots) cluster together although two of the non-pregnant (non-PR) cows (green dots) cluster together with the PR group. Numbers in parentheses correspond to the animals’ IDs.

### Functional analysis

- **Gene set enrichment analysis (GSEA):** GSEA was applied to identify significantly enriched pathways considering the entire expression pattern change between PR and non-PR cows. The functional enrichment with GSEA determined 11 pathways enriched of genes up-regulated in PR cows (Supplementary Table 2A) and four pathways for genes down-regulated in PR cows (Supplementary Table 2B).
- **Gene ontology (GO) term analysis and functional classification:** The NETwork-based human gene enrichment (NET-GE) GO term enrichment analysis performed on all DEG (up-and down-regulated) identified a significant enrichment for two cellular component (CC) GO terms (cell surface and extracellular matrix part), two molecular function (MF) GO terms (Leukotriene-C4 synthase activity, Type I TGF-β receptor binding) and 41 BP GO terms (Supplementary Table 3). The BP GO terms were mainly related to response to growth factor, negative regulation of TGF-β signaling, extracellular matrix organization and cell adhesion proliferation and migration as well as to arachidonic acids and eicosanoid metabolic process and immune response and leukocyte proliferation. Other interesting GO terms were response to progesterone and estrogen. The functional classification provided by IPA® revealed involvement of cellular processes (cellular, function, movement, morphology, development, growth and proliferation free radical scavenging) as well as the importance of cellular metabolism of lipids, carbohydrate, vitamins and amino acids (Supplementary Table 4).
- **Up-stream regulator analysis with Ingenuity^®^ Pathway Analysis (IPA^®^)**: We selected upstream regulators that were predicted as significantly activated after bias correction by IPA®. We identified 11 as being activated and two as being inhibited in PR cows (Table 1). The top 25 identified upstream regulators are listed in Supplementary Table 5.

**Table 1.**
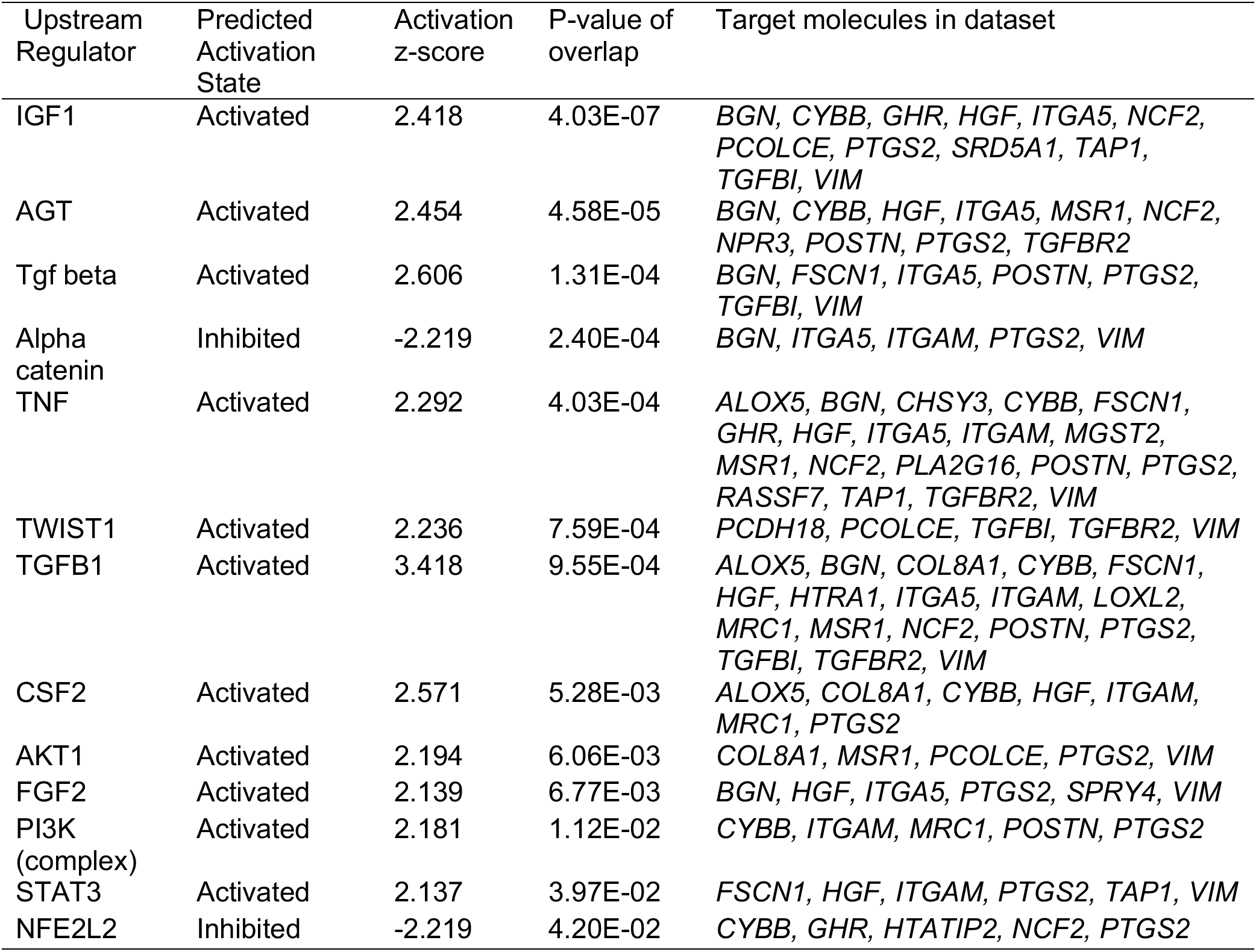
Up-stream regulators predicted as being activated or inhibited in pregnant cows compared to non-pregnant cows. Upstream regulator with significant Z score (|Z-score| >2) and P value of overlap (P value< 0.05) are listed ordered by P-value of overlap.

Integration of the results from the functional analysis unveiled five main biological mechanisms associated with endometrial receptivity, shown in detail in Fig. 2. These mechanisms are: a) Extracellular Matrix (ECM) interaction (Fig. 2A), b) control of histotroph metabolic composition (Fig. 2B), c) prostaglandins (PG) synthesis (Fig. 2C), d) TGF-β signaling (Fig. 2D) and e) inflammation and leukocyte activation (Fig. 2E). In addition, secondary pathways and upstream regulators previously associated with endometrial receptivity were identified (Fig. 2i-iii).

**Figure 2.**
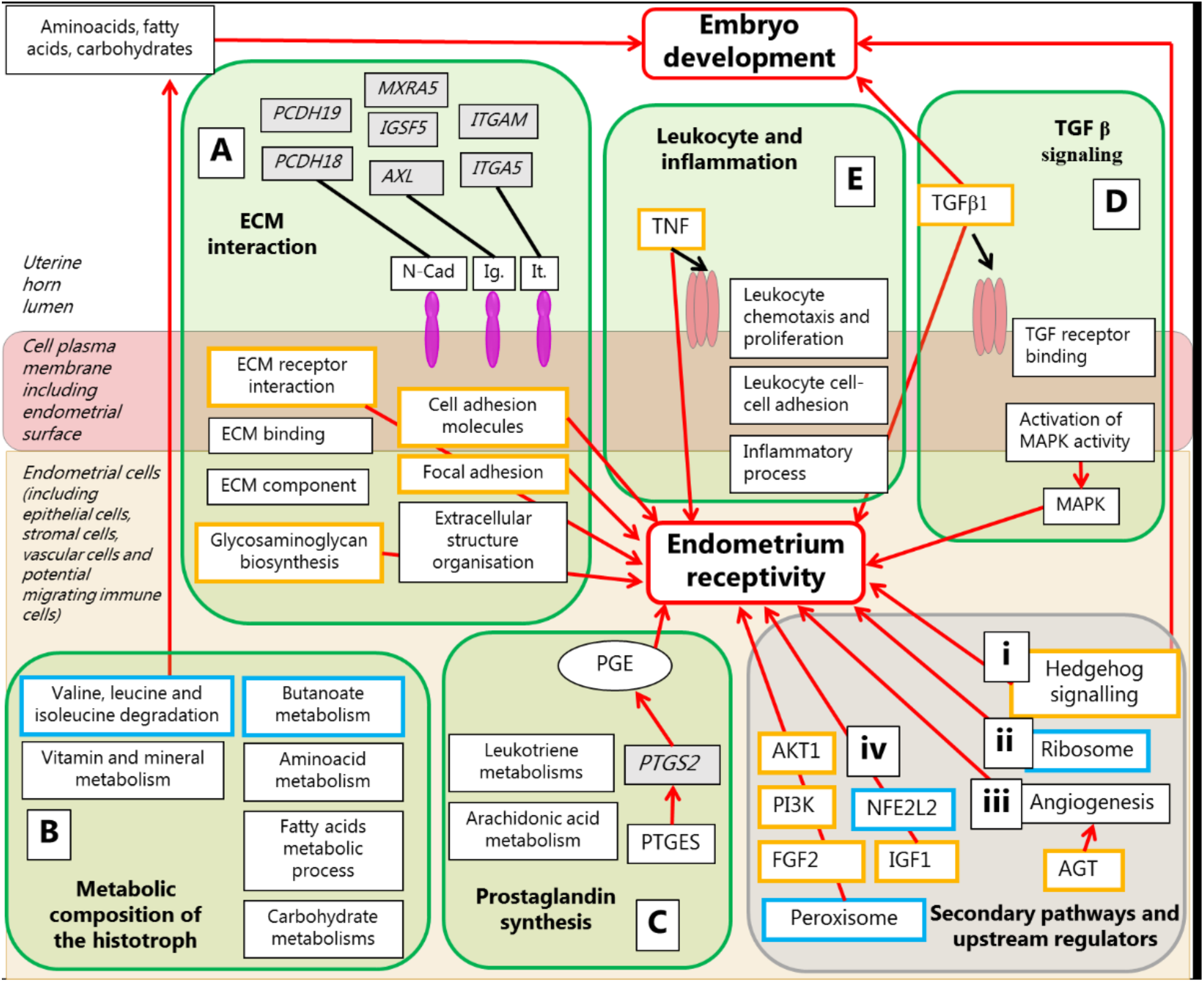
Summary of the biological mechanisms, upstream regulators and genes with their activation state in pregnant (PR) cows and their localization in the uterus. Localization in the uterus: uterine horn lumen (white), cell plasma membrane including endometrial surface (purple) and endometrial cells including epithelial cells, stromal cells, vascular cells and potential migrating immune cells (pink). Rounded shaped rectangles in green = five main mechanisms identified: (A) ECM interaction, (B) metabolic composition of the histotroph, (C) prostaglandin synthesis, (D) TGF β signaling, and upstream regulators (E) leukocyte and inflammation. Rounded shaped rectangle in grey = secondary pathways and upstream regulators: (i) Hedgehog signaling, (ii) ribosome, (iii) angiogenesis, (iv) other upstream regulators. Grey filled rectangles = genes up-regulated in PR cows. Rectangles = upstream regulators and biological processes predicted to be up regulated-regulated (orange) or down-regulated (blue) in PR cows or with no specific direction (black). Ovals = other molecules from literature. The discontinuous lines point to the biological functions, genes and upstream regulators belonging to the five main mechanism. Red arrows = positive effect exerted on PR endometrium (embryo development and endometrial receptivity). The black lines points to the three type of cell adhesion molecules (CAD) colored in purple (N-cad = N- cadherins, Ig = immunoglobulins, It = integrins) that are encoded by the up-regulated genes. The black arrows show the mechanisms of TNF and TGFb, which bind to the correspondent receptors (pink molecules) in the endometrial surface.

### Comparison with other studies

Results from this analysis are shown in Fig. 3A. There was a significant association between overlapping genes going in the same direction between PR and cows with good genetic merit for fertility, or high-fertile (HF) cows (when compared, respectively, to non-PR and low-fertile cows). On the contrary, some genes overlapped in opposite direction between PR and cows with high levels of progesterone (HP) (versus their respective controls) but there was no significant association. The multidimensional scaling (MDS) plot (Fig. 3B) depicts the distribution of PR and non-PR samples based on the expression of the overlapping genes. Samples were distributed apart when the expression of the overlapping genes with HF were considered, but not with the expression of the overlapping genes with HP.

**Figure 3.**
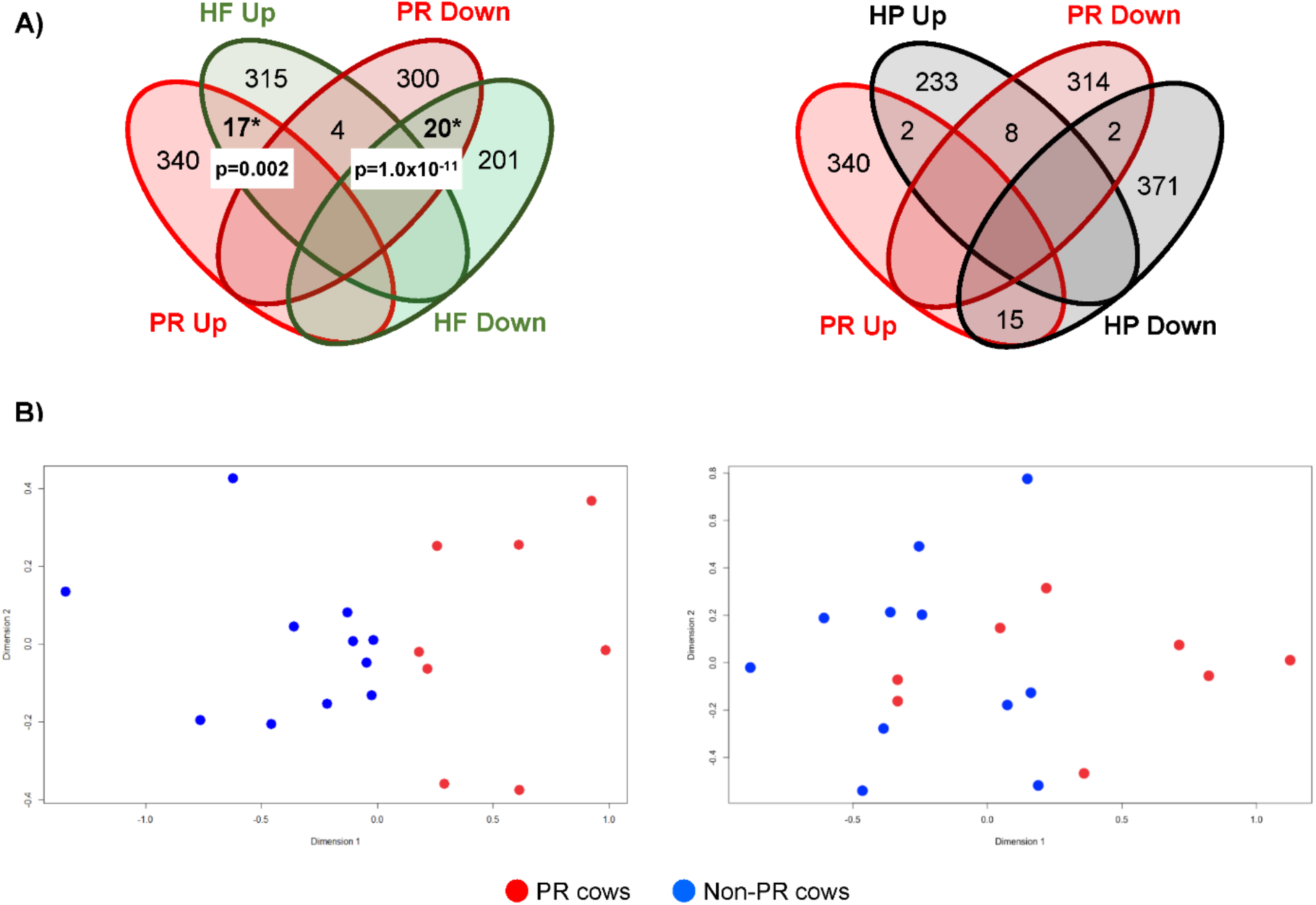
Comparison of differentially expressed genes (DEG) in the endometria of pregnant (PR) cows and DEG in the endometria of high fertility (HF) cows or cows treated with a progesterone releasing device (HP); relative to their respective control groups. (A) The Venn diagrams show the number of up- and down-regulated genes for each comparison, the number of overlapping genes in the same or opposite direction and the corresponding significant p-value. (B) The multidimensional scaling plots display the distribution of the PR (red dots) or non-pregnant (non-PR) cows (blue dots) according the expression of the overlapping genes with the HF group (left plot) or HP group (right plot).

These results reinforce the concept that the observed differences in the endometrial transcriptome between PR and non-PR cows is given due to the inherent molecular signature for each cow, rather than to variations in hormonal concentrations.

## Discussion

Understanding the physiology of the bovine endometrium and its role in the establishment of pregnancy has been an overarching theme in cattle reproduction. Transcriptomic analyses have become a powerful tool for obtaining in depth molecular knowledge about endometrial function. Exploration of the transcriptome provides an unbiased system-wide analysis of physiologically relevant gene expression levels, determining endometrial receptivity (reviewed by Forde & Lonergan ^26^). It has been demonstrated that the endometrium is already modified at day 5-6 after ovulation, independently of the presence of an embryo, in order to create a micro-environment to receive the embryo from the oviduct ^26^. Interestingly, results from this and other studies indicate that the endometrial transcriptome at day 7 of animals that become pregnant differs from those that do not ^19,21,22^. Therefore, this period is ideal to determine the receptivity of a cow, which is particularly relevant in ET procedures when highly valuable embryos are transferred.

Previous investigators have explored the endometrial transcriptomic profile during this period of the estrous cycle. Slilew-Wondim et al. ^22^ collected endometrial samples at days 7 and 14 day while Ponsuksili et al. ^19^ did it at days 3 and 7. Both studies implemented a similar model used in the present investigation, in the sense that *in-vivo* produced or IVP embryos, respectively, were transferred in the subsequent cycle and animals were retrospectively classified according to their pregnancy status. However, these studies were carried out in Simmental heifers and the biopsies were obtained using the cytobrush technique, which mainly yields cells from the endometrial epithelium. In another study, Binelli et al. ^21^ inseminated Nellore cows and collected endometrial samples 6 days later, from the uterine horn contralateral to ovulation. Cows were also classified retrospectively according pregnancy diagnosis. On a different approach, Killen et al. ^27^ classified beef heifers *a priori,* based on the pregnancy results obtained after 4 rounds of AI. Samples were obtained at day 7 from the intercaruncular sites of the uterus after animals’ slaughter. Finally, Moran et al. ^24^ collected endometrial samples trough biopsies in day 7 of the cycle from lactating dairy cows, similar to the present study. Nevertheless, they classified the animals based on the genetic merit for fertility, and thus not on the pregnancy status. All these studies used different synchronization methods to induce estrous.

Results from these studies highly contribute to the knowledge of the endometrial transcriptomic profile at the time when an embryo is transferred into a recipient. However, they also demonstrate that the endometrial transcriptome is affected by many factors, including parity (heifers or cows), breeds (dual-purpose, beef or dairy breed), and induction of estrus ^17^.

Establishment of pregnancy is dependent on a relevant transcriptome conveying endometrial receptivity, but also on embryonic viability. As alluded to earlier, a multitude of factors influence embryo viability; a matter of particular importance for IVP embryos. Here, we aimed at minimizing these sources of variation as much as possible by e.g. using a single bull, a consistent IVP system including extensive control material, a single person performing the ET as well as careful matching of day of transfer and stage of embryonic development.

In the present study, lactating dairy cows were observed and monitored for spontaneous estrus, and hence, no hormonal synchronization method was applied. Cows were classified retrospectively according to pregnancy status of IVP embryos transferred on the next cycle after the endometrial biopsies were obtained, which were done with an instrument to include the full endometrium (luminal epithelium, glandular epithelium, stroma, blood vessels, nerves and invading leukocytes). Also, biopsies were taken consistently from the cranial to middle part of the uterine horn ipsilateral to the CL, in order to investigate specifically the part that receives the embryo at ET and to avoid regional differences in the uterine transcriptome ^20^. Therefore, our findings could be of enormous help to identify key genes that characterize a receptive endometrium. Results from this and other studies could be used to predict endometrial receptivity of a cow, as it has been developed for humans ^28^, ideally through the development of measurable biomarkers from accessible samples, such as the blood.

Results from differential gene expression between PR and non-PR cows revealed 111 candidate genes, with 60 up-regulated and 51 down-regulated in PR cows. Furthermore, the functional analysis of these DEG identified five main biological mechanisms associated with endometrial receptivity (depicted in Fig. 2): a) ECM interaction, b) control of histotroph metabolic composition, c) PG synthesis), d) TGF-β signaling and e) inflammation and leukocyte activation. Each of these mechanisms is briefly discussed in the following paragraphs.

a. Extracellular matrix interaction: Involvement of many DEG (up-regulated in PR cows) in the ECM was confirmed by the enrichment for MF GO terms (glycosaminoglycan binding) and for the BP GO term (regulation of adhesion, ECM organization, extracellular structure organization). The GO terms belonging to the CC domain revealed that a huge component of the proteins encoded by the DE genes are secreted or membrane bound, and that many of them are part of the ECM. Among them, 16 proteins showed to function as anchoring junctions important for the endometrial organization. In human, three super-families of proteins are involved in ECM adhesion and embryo implantation ^29^, which were part of the DEGs: three immunoglobulins, two cadherins and two integrins. These findings reflect that the endometrial tissue is rich in ECM molecules for interaction and adhesion. A balanced ECM integrity has been associated with pregnancy maintenance, and the combined action of metalloproteinase inhibitor (TIMP2) and metalloproteinases ^30^ has been suggested to be responsible for the regulation of ECM integrity ^30,31^. In addition, genes related with the ECM have been also deregulated in several of the studies mentioned above ^21,22,24^.
b. Histotroph metabolic composition: The histotroph is the mixture of molecules secreted into the uterine lumen and, thus, its composition cannot be determined through an endometrial biopsy. However, the endometrial transcriptome reflects pathways that are involved in the cellular production of components of the histotroph. Our transcriptomic results demonstrated alterations related to the histotroph synthesis that could be involved in increased receptivity in the PR cows. A set of genes down-regulated in PR cows was enriched for the KEGG pathway valine, leucine and isoleucine degradation. Amino acids are important components of the histotroph ^32^ and are essential for conceptus elongation ^33–35^. Bovine embryos utilize amino acids ^34,36^, and the concentrations of L-leucine, L-valine and L-isoleucine increase in pregnant animals ^34,37^. We speculate that the early inhibition of endometrial genes involved in degradation of these amino acids would lead to a higher concentration of them during the implantation window, improving the endometrial receptivity. In addition, some of the DEG were involved in molecular transport and in the metabolic pathways of carbohydrates, vitamins, minerals and amino acids, according the IPA® classification. Again, these are important components of the histotroph that show how PR and non-PR cows differ regarding the quality of this essential nourishment for the embryo ^31,38^.
c. PG synthesis: The GO term analysis revealed a set of genes involved in arachidonic acid metabolic processes and leukotriene C4 synthases. We identified PTGES as the upstream regulator of four DEG (CYBB, EZR, PTGS2 and VIM). PTGES is an enzyme that generates PGE from PGH2 ^39^, which is derived from an intermediate step performed by PTGS2. Interestingly, PTGS2 was differentially up-regulated in PR cows. In sheep, PGs generated by the PTGS2 in the conceptus were responsible for regulation of genes involved in conceptus elongation ^33^.
d. TGF-β signaling: One of the TGFβ signaling pathways is the MAPK cascade that was one of the biological functions enriched with the DEG list (both up- and down-regulated in PR cows). TGFβ1 and its related molecules were identified as upstream regulators of the DEG, and their encoded proteins were predicted to be activated in PR cows. Furthermore, the TGFβI and TGFBR2 were up-regulated in PR cows. The TGF-βs control many cellular processes and the mechanism is exerted through interaction with the ECM. This signaling pathway stimulates ECM formation and prevents its degradation ^40,41^. Genes belonging to the TGF-β pathway regulate pre-implantation development in both embryo and endometrium ^42^. Furthermore, TGF-β1 and TGF-β2 stimulate proliferation of bovine trophoblast and fetal endometrial epithelial cell in-vitro ^43,44^. Therefore, activation of the TGF signaling at diestrus could lead to a better embryo receptivity, supporting embryo development and elongation and at the same time stimulating ECM formation.
e. Inflammation and leukocyte activation: The DEG enriched for inflammation response and for leukocyte chemotaxis, adhesion, and proliferation. In cattle, leukocyte presence seems not to be a requirement for placentation ^45,46^ as in humans ^47^. However, the innate immune system is activated during early pregnancy in the bovine uterus ^48^ and several chemokines known to affect immune tolerance (i.e. leukocytes) are up-regulated in pregnant animals. These actions are required for establishment of the right balance between pro-and anti-inflammatory molecules during initial placentation ^48^. Immune related genes were found also to be deregulated in the endometrial samples from day 6-7 of the estrus cycle in most of the studies mentioned at the beginning of this section ^19,22,24,27^.

In addition, secondary pathways and upstream regulators previously associated with endometrial receptivity were identified, such as: (i) The Hedgehog pathway (Fig. 2i), constituted by genes up-regulated in PR cows, involved in implantation and early embryo development ^49,50^. (ii) The ribosome KEGG pathways (Fig. 2ii), sustained by genes down-regulated in PR cows, is likewise downregulated in the endometrium during diestrus ^51^ corresponding to the phase of initial embryonic development and adhesion, and (iii) The angiogenesis process (Fig. 2iii), including the upstream regulator predicted to be activated in PR cows (AGT), which contributes positively to the establishment of endometrium receptivity ^19,52^.

The upstream regulators of the DEG were explored in the present study as well. Progesterone was identified as the top upstream regulator of the DEG, being associated with the profile of 15 genes. Moreover, response to progesterone and estrogen were enriched processes among the DEG. Lower progesterone concentration has a negative effect on expression of genes that regulate histotroph composition and conceptus elongation ^18,26^. The upstream regulator analysis did not predict any significant accumulation or depletion of progesterone from the analysis of gene expression changes of its target molecules. However, 11 upstream regulators were predicted to be activated and two to be inhibited in PR cows. Among them, IGF1 ^27,53,54^, AGT ^52,55^, TNF ^56,57^, AKT and PI3K ^58,59^, FGF2 ^20^, TGFβ1 (previously discussed) (activated in PR cows) and NFE2L2 ^19^ (inhibited in PR cows) have been previously associated with endometrial processes and receptivity.

Given the central role of progesterone and estrogen in defining endometrial receptivity, an additional analysis was performed to characterize the molecular profile of PR and non-PR cows. Two external datasets, obtained from public domains, were re-analyzed in order to compare the corresponding lists of DEG with the ones from the present study. One of the datasets, introduced earlier on this section, corresponded to endometrial samples obtained at day 7 from lactating dairy cows with good (HF) or poor (LF) genetic merit for fertility. Cows classified as HF presented greater body condition score and circulating IGF1 concentrations throughout lactation, more favorable metabolic status during the transition period and faster uterine recovery and earlier resumption of cyclicity after parturition ^24^. The other dataset corresponded to a large study with several experiments where samples were obtained from pregnant or cyclic cows at different time points in the cycle ^18,25,60^. For the present study, only samples collected at day 7 from cyclic heifers with high (HP) or normal levels of progesterone were considered. Animals in the HP group were treated with a progesterone releasing intravaginal device (PRID) from day 3 of the estrus cycle ^25^. Thus, that experiment aimed to identify the effects of elevated progesterone levels on the endometrial transcriptome.

Comparison of the up and down regulated DEG on the present study with the DEG from these two studies, showed a significant association of genes going in the same direction for PR and HF cows, while no association was observed with DEG in HP cows (Fig. 3). These results suggest that the endometrial transcriptome of PR cows is more similar to the one from HF cows than to the one influenced by high progesterone levels. Thus, inherent characteristics of PR and non-PR cows would be defining the differences in endometrial transcriptome rather than hormonal variations between these two groups.

## Conclusion

Findings from this study provide knowledge about the endometrial molecular signature required for pregnancy establishment, around the time of embryo transfer, in lactating dairy cows. Five main relevant mechanisms were identified: ECM interaction, histotroph metabolic composition, prostaglandin synthesis, TGF-β signaling and inflammation and leukocyte activation; all of which appear to be regulated by progesterone and estrogen concentrations. Moreover, we identified candidate up-stream regulators that were predicted to be activated (IGF1, AGT, TNF, AKT, PI3K, FGF2, TGFβ1) or inhibited (NFE2L2) in PR cows. In addition to our own experimental datasets, we analyzed two public gene expression datasets from similar experiments and compared up-and down-regulated DEG from our study with the DEG from these two studies. Results confirmed significant association of genes going in the same direction for high receptive cows. Hence, these candidate gene regulators and enriched pathways could be used for identification of potential biomarkers for predicting pregnancy success in IVP-ET in artificial reproduction technology. However, further experimental validation in large samples is recommended.

## Material and methods

### Animals

This study was carried out in strict accordance with the Danish law on animal experimental work (law no. 474 of 15 May 2014). According to this law, an application to perform the live animal work in this study was submitted and approved by the ethical committee “Danish Animal Experiments Inspectorate” before starting the experiment (license no. 2015-15-0201-00636). The animals were Holstein Friesian cows in first, second or third lactation, with no clinical disease and no history of reproductive problems. All estruses were spontaneous during the experimental period, so no hormonal synchronizations were used. Estrus observation was performed visually twice daily and by automatic monitoring of cow activity in 6-h intervals. Endometrial biopsies were taken from cows on day 6-8 in diestrus (day 0 = last day of standing estrus) followed by ET on day 6-8 in the following cycle. The pregnancy status was determined after slaughter on day 26-47.

Embryos were produced in two or three replicates per week in order to ensure availability of fresh embryos for transfer according to the recipient’s natural cycle (3500 oocytes fertilized with semen from one bull in 35 IVP rounds). For transfer, 28 blastocysts were selected, derived from 16 IVP replicates (1600 oocytes fertilized) with an average blastocyst rate of 43% (min-max 26-56%).

The experiment was continued until 12 pregnancies were achieved in order to obtain enough replicates to get reliable results with DESeq2 ^61^. The overall pregnancy rate obtained was 43% using a total of 28 cows for ET. From the 16 non-pregnant cows sampled, 12 were selected for further analysis to obtain balanced experimental groups (considering parity number). Among the 24 cows selected, both groups were balanced regarding parities: in the PR group, seven cows had one calving, two cows had two calvings and three cows had three calvings, while in the non-PR group, five cows had one calving, three cows had two calvings and four cows had three calvings. The 24 cows had an average age at biopsy collection of 1208 days (SD = 388, min = 768, max = 1789). The whole experiment took around 6 months.

### Endometrial biopsies

On day 6-8 of the cycle preceding embryo transfer, the cows had an epidural analgesia (5 ml Procamidor, Salfarm Danmark A/S, Denmark), and an endometrial biopsy of 50-60 mg was taken from the middle to cranial part of the uterine horn ipsilateral to the CL using a biopsy instrument (141700, Kruuse A/S, Denmark). The biopsies were transferred into Eppendorf Safe-Lock Microtube 2.0 ml with 1.5 ml RNAlater (76154, Qiagen), and kept at room temperature for minimum two hours. Each biopsy was divided into two replicates (each of 25-30 mg tissue), and transferred into 0.5 ml RNAse-free Microfuge tubes (Ambion) and stored at −80ÁC.

The biopsies were taken from live cows. Therefore, they could not be taken precisely in relation to the caruncle sites. The biopsies were deep enough to include the full endometrium (luminal epithelium, glandular epithelium, stroma, blood vessels, nerves and invading leukocytes). Exclusively endometrial biopsies of the expected weight, i.e. (50 mg) were included in the full bioinformatic analyses in order to achieve a comparable uniform material. Thus, 2 samples from PR cows were excluded from the RNA-Seq analysis.

### RNA extraction and sequencing

Endometrial biopsies were homogenized with Tissue Lyser LT. Total RNA was isolated and purified with RNAeasy mini Kit. Quantity and quality of the RNA samples were assessed by Agilent 2100 Bioanalyzer. One sample was not sequenced because its RNA integrity number, that is a measure of the RNA quality, was 1.5, resulting in 11 samples from non-PR cows.

The libraries for the sequencing were generated with an Illumina TruSeq-stranded total RNA protocol. All RNA-Seq data have been deposited in NCBI’s Gene Expression Omnibus and are accessible through GEO accession number GSE115756. (https://www.ncbi.nlm.nih.gov/geo/query/acc.cgi?acc=GSE115756).

RNA samples were paired-end sequenced with the Illumina HiSeq 2500 platform with a read length of 100 nt, in two lanes pooling all the samples together.

### In-vitro production of embryos and embryo transfer

All chemicals were purchased from Sigma–Aldrich Corp. (St Louis, MO, USA) except otherwise indicated. For composition of media (sperm-TALP, IVM, IVF and IVC), see Supplementary Table 6. A detailed description of the IVP protocol is found in Holm et al. ^62^.

On day 6-8, each recipient cow had one fresh embryo transferred. On the transfer day (day 7 of embryo development, IVF = day 0), the embryos were evaluated and selected based on developmental stage and morphology. Morphological good and excellent blastocysts/expanded blastocysts were used for transfer. For synchronization between embryo development stage and recipient, day 6 recipients had a blastocyst transferred while day 7 and 8 recipients had a blastocyst or an expanded blastocyst transferred. Before transfer, the embryos were washed in holding medium (Syngro Holding, Bioniche), loaded into 0.25 ml straws and transported to the stable at 38°C. The cows had an epidural analgesia (5 ml Procamidor) before the one embryo was transferred to the middle to cranial part of the uterine horn ipsilateral to the CL.

### Samples at slaughter

From day of transfer until day 26-47, the recipient cows had not shown any signs of estrus and were then slaughtered to collect their reproductive organs. The cows were slaughtered across different days to describe gradual changes in pregnancies based on IVP embryos, which were destined to another experiment. After slaughter, the uterine horns were opened, and based on the findings of embryos(26-42 days of age), fetuses (above 42 days of age), and fetal membranes, the cows were registered as pregnant (PR) or non-pregnant (non-PR). By this approach, we expected to find signs of an eventual early loss of an early pregnancy, because of the size of an early embryo with membranes, and because the uterus had not been cleaned by an estrus period.

### Bioinformatic analysis

#### RNA processing

The quality of the reads was checked with *FastQC* (v. 0.11.2) ^63^. Adapters and quality trimmer and filtering were performed using *Trimmomatic* (v. 0.36) ^64^ to compute (1) trimming of the 3’-end and 5’-end with a quality threshold of 20, (2) window based filtering using a window of 4 bases and an average quality threshold value of 15, (3) filtering reads with a final length of less than 25 nucleotides. The remaining 41,799,192 read pairs were mapped to the bovine reference genome (*Bos taurus* UMD3.1) ^65^ with *STAR aligner* (v. 2.5.2) ^66^, including the gene *Bos taurus* release 87 annotation in the genome index. We allowed for a maximum of five mismatches, while setting the other parameters to *STAR* default values. Read counts were estimated at gene level using *HTSeq-count* (v. 0.6.0) ^67^, setting the model of intersection as ñintersection-nonemptyî using the same annotation file.

#### RNA-Seq preprocessing

The samples 6 and 23, from PR cows, showed different profiles during library fragmentation and were excluded from the rest of the analysis (Supplementary Fig. 2). Further quality control on count data were performed with *Noiseq* (v.2.14.0) ^68^ and with the R package cqn ^69^. The quality control did not reveal any GC or length bias between samples (Supplementary Fig. 3).

#### Differential gene expression analysis

We filtered out genes with less than one count per million in more than eight samples (smaller class) ^70^. The dataset used in the differential expression analysis consisted of 19 samples (eight pregnant and 11 non-pregnant) and 14231 genes. The gene expression counts were normalized by library size with DESeq2 methods.

The differential gene expression analysis was performed by fitting a logistic regression model to the gene counts (modelled by a negative binomial distribution) in DESeq2 as:

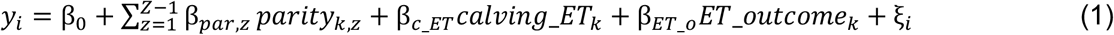

Where y_i_ is the gene normalized gene expression count for gene *i*, β_0_ is a fixed intercept term; parity i_(z,*f*Z−1)_ is a set of (Z−1) dummy variables representing the parity for the *i*^th^ animal; and calving, ET representing the days between the last calving and the embryo transfer was fitted as covariate. Β_parity,z_, and β_c_ET_ are the corresponding solutions. The differential expression analysis was based on the embryo transfer outcome that was fitted as a dummy variable (0=non-pregnant or 1=pregnant). We applied a Wald test statistic ^61^ to test for model significance. P values were adjusted with Benjamini-Hochberg method. Differentially expressed genes were called at False Discovery Rate (FDR) lower than 0.05.

#### Functional enrichment analysis

The GSEA was performed using the *GseaPreranked* tool from GSEA v2.2.2 (Broad Institute) ^71^. This tool tests functionally related set of genes which are defined *a priori*. The DEGs overlapping with the gene sets that are found at the top or bottom of the list are those that contribute the most to the enrichment results, these genes define the core enrichment. The enrichment score is computed using a weighted Kolmogorov ‒Smirnov-like statistic. The normalized enrichment score is computed to account for sizes of the gene sets. We used the gene sets from Molecular Signatures Database (MSigDB Collections) v5.1. GSEA was able to map 12,391 genes based on the gene symbol.

The functional analysis with using IPA (QIAGEN, CA, USA) (www.qiagen.com/ingenuity) was performed using all the DEG, using the Entrez ID as identified. IPA® mapped successfully 99 out of the 111 DEG. IPA® defines upstream regulators as any type of molecules that could affect the expression of a specific set of genes. The upstream regulator which are significantly overrepresented for the DEG list, are selected. If the directionality of the gene expression (up-regulation, down-regulation) of the DEGs is consistent with the a priori information of that upstream regulator, an additional information about its predicted status (activated or inhibited) is provided.

The proportions of the DEG were not correlated to the length of the genes (Supplementary Fig. 4). Thus, the GO term enrichment analysis was performed with NET-GE ^72^, using the symbol name for Biological Process (BP), Cellular Component (CC) and Molecular Function (MF) GO terms. NET-GE is network-based method for gene enrichment analysis. For each functional gene set NET-GE relies on an interactome to build a module of functionally related genes. The modules are then tested via Fisher’s exact test.

Results of the functional analysis were considered statistically significant with a FDR lower than 5%.

### Comparison with other studies

As a complement of our study, two publicly available gene expression dataset were used to compare with our results. These datasets were downloaded from the NCBI GEO repository, with the accession numbers GSE52438 ^24^ and GSE33030 ^25^. In the first study (GSE52438), endometrial biopsy samples from non-pregnant lactating dairy cows on day 7 of the estrus cycle with similar genetic merit for milk production traits but with very good genetic merit for fertility (HF, n=7) and cows with very poor genetic merit for fertility (LF, n=7) were analyzed by RNAseq technology. The matrix of gene counts was downloaded and re-analyzed with the DESeq2 package for R ^61^.. Genes with low expression counts (less than 1 CPM) in 7 or more samples, were filtered out before normalization. Normalization method by library size was performed according DESeq2 methods. Differentially expressed genes were determined using a generalized linear model where the counts for each gene on each sample are modeled using a negative binomial distribution with fitted mean and a gene-specific dispersion parameters. The Wald test statistic is applied to test for significance of the generalized linear model coefficients. In the second study (GSE33030), samples were obtained from pregnant or cyclic cows at different time points in the cycle ^18,25,60^ but only samples collected at day 7 from cyclic heifers were considered here. Animals were treated with a progesterone releasing intravaginal device (PRID) from day 3 of the estrus cycle (HP group, n=5) or did not receive a device (NP group, n=5). Extracted RNAs were hybridized to the Affymetrix Bovine Genome Array platform. The raw data was obtained and processed with the gcRMA package ^73^, which was used to import the raw data into R, perform background correction, and then transform and normalize the data using the quantile normalization method. DEG were determined with moderated t-statistics, a variation of the classical t-test ^74^.

For the sake of comparisons, significant DEGs in these and our study were defined as those with a p-value < 0.05 and a fold change > 1.4. Up or down-regulated genes in our study were compared to the up-and down-regulated genes in the two external studies. The ENSENMBL IDs were used for comparisons. The test of independence (Pearson’s chi-square test) was employed to determine the relatedness of up-and down-regulated DEGs observed in PR vs non-PR with up-and down-regulated DEGs in HF vs LF or HP vs NP. Finally, similarities between PR and non-PR samples according to the expression of the overlapping genes was assessed trough a multidimensional scaling plot (MDS), performed with the Glimma package ^75^.

## Acknowledgements

Authors thank technicians involved in experimental work.

## Availability of data

All data are fully available without restriction on NCBI GEO database https://www.ncbi.nlm.nih.gov/geo/query/acc.cgi?acc=GSE115756

## Authors’ contribution

HNK first conceived the design of this study, the GIFT (Genomic Improvement of Fertilization Traits) research project, managed the project and contributed to design of bioinformatics and statistical analyses; PH and HC contributed to further development of the design and experiments. HSP and HC conducted the IVP-ET experiments. GM analyzed RNA-Seq transcriptomic data and interpreted the results. MBR contributed to the analysis and further improvement in writing and interpretation. GM, HSP and MBR wrote the first draft of this manuscript. All authors contributed to the writing of this manuscript and approved the final version.

## Ethical Statement

This study was carried out in strict accordance with the Danish law on animal experimental work (law no. 474 of 15 May 2014). According to this law, an application to perform the live animal work in this study was submitted and approved by the ethical committee ÒDanish Animal Experiments InspectorateÓ before starting the experiment (license no. 2015-15-0201-00636).

