## Supplementary for "Characterization of the Endometrial Transcriptome in Early Diestrus Influencing Pregnancy Establishment in Dairy Cattle after Transfer of In-Vitro Produced Embryos"

**Supplementary Tables: SM1, SM2, SM3, SM4, SM5, SM6 and SM7**

**Supplementary Figures: SM1, SM2, SM3 and SM4**

**Supplementary Tables**

***Table SM 1. List of differentially expressed genes between pregnant and non-pregnant cows***

| Gene ID | Gene Name | Description | FC | P-adj | IPA classification |
|---|---|---|---|---|---|
| ENSBTAG00000000575 | *TNC* | Tenascin C | 2.51 | 7.91E-05 | Not annotated |
| ENSBTAG00000011381 | *SLC30A3* | Zinc transporter 3 | 0.47 | 1.13E-02 | Transmembrane receptor |
| ENSBTAG00000017677 | *SCG3* | Secretogranin-3 | 1.96 | 2.80E-02 | Enzyme |

| | | | | | |
|---|---|---|---|---|---|
| ENSBTAG00000013662 | *COL8A1* | Collagen type VIII alpha 1 chain | 1.96 | 2.80E-02 | Other |
| ENSBTAG00000009513 | *TGFBI* | Transforming growth factor-beta-induced protein ig-h3 precursor | 1.95 | 7.50E-03 | Other |
| ENSBTAG00000013990 | *WNT6* | Wnt family member 6 | 1.91 | 3.06E-02 | Other |
| ENSBTAG00000008708 | *GPC5* | Glypican 5 | 0.53 | 3.27E-02 | Other |
| ENSBTAG00000033153 | *GRIK2* | Glutamate receptor, ionotropic kainate 2 | 1.89 | 1.88E-02 | Other |
| ENSBTAG00000014127 | *PTGS2* | Prostaglandin-endoperoxide synthase 2 | 1.88 | 3.14E-02 | Enzyme |
| ENSBTAG00000048257 | *CYP4F3 (Uniprot)* | Cytochrome P450, family 4, subfamily F, polypeptide 3 | 0.54 | 4.01E-02 | Other |
| ENSBTAG00000002092 | *PI16* | Peptidase inhibitor 16 | 0.54 | 3.92E-02 | Transporter |
| ENSBTAG00000027320 | *KCNB1* | Potassium voltage-gated channel subfamily B member 1 | 0.54 | 2.80E-02 | Other |
| ENSBTAG00000017020 | *S100G* | Protein S100-G | 0.55 | 2.80E-02 | Other (cadherin) |
| ENSBTAG00000002444 | *MKI67* | Marker of proliferation Ki-67 | 1.83 | 3.92E-02 | Enzyme |
| ENSBTAG00000000198 | *LYZ (homology )* | Lysozyme | 0.55 | 4.26E-02 | Ion channel |
| ENSBTAG00000003920 | *TGM1* | Transglutaminase 1 | 0.55 | 4.98E-02 | Enzyme |
| ENSBTAG00000007665 | *NPR3* | Natriuretic peptide receptor 3 | 1.80 | 4.98E-02 | Not annotated |
| ENSBTAG00000009226 | *SCUBE3* | Signal peptide, CUB domain and EGF like domain containing 3 | 1.79 | 4.01E-02 | Other |

| | | | | | |
|---|---|---|---|---|---|
| ENSBTAG00000008255 | *QRFPR* | Pyroglutamylated rfamide peptide receptor | 1.79 | 4.14E-02 | Enzyme |
| ENSBTAG00000014169 | *PCDH19* | Protocadherin 19 | 1.78 | 1.82E-02 | Other (cadherin) |
| ENSBTAG00000009230 | *FBLN7* | Fibulin 7 | 1.78 | 3.20E-02 | Enzyme |
| ENSBTAG00000006214 | *LOXL2* | Lysyl oxidase homolog 2 precursor | 1.75 | 2.33E-02 | Other |
| ENSBTAG00000012815 | *CLEC12A* | C-type lectin domain family 12 member A | 1.75 | 1.28E-02 | Enzyme |
| ENSBTAG00000015749 | *STEAP1* | STEAP family member 1 | 1.74 | 4.98E-02 | Not annotated (immunoglobulin) |
| ENSBTAG00000010986 | *KCNQ1 (homology)* | KCNQ1potassium voltage-gated channel subfamily Q member 1 | 0.58 | 3.81E-02 | Other |
| ENSBTAG00000022150 | *MXRA5* | Matrix remodeling associated 5 | 1.73 | 3.69E-02 | Ion channel |
| ENSBTAG00000019761 | *MANBA* | Mannosidase beta | 0.58 | 3.28E-02 | Phospatase |
| ENSBTAG00000031503 | *NDUFA4L2* | NADH dehydrogenase | 1.73 | 3.89E-02 | Not annotated |
| ENSBTAG00000012718 | *XK* | X-linked Kx blood group | 0.58 | 1.88E-02 | Other |
| ENSBTAG00000014661 | *CHSY3* | Chondroitin sulfate synthase 3 | 1.71 | 4.26E-02 | Peptidase |
| ENSBTAG00000003671 | *SOWAHA* | Sosondowah ankyrin repeat domain family member A | 0.59 | 2.41E-02 | Other |
| ENSBTAG00000020652 | *TSSK3* | Testis-specific serine/threonine-protein kinase 3 | 0.59 | 3.90E-02 | Other |
| ENSBTAG00000022028 | *DERL3* | Derlin 3 | 0.59 | 4.01E-02 | Enzyme |

| | | | | | |
|---|---|---|---|---|---|
| ENSBTAG00000009076 | *ADD2* | Adducin 2 | 1.70 | 3.81E-02 | Enzyme |

| | | | | | |
|---|---|---|---|---|---|
| ENSBTAG00000020319 | *ALOX5* | Arachidonate 5-lipoxygenase | 1.67 | 3.06E-02 | Other |
| ENSBTAG00000012409 | *POSTN* | Periostin | 1.66 | 4.26E-02 | G-protein coupled receptor |
| ENSBTAG00000011545 | *GDA* | Guanine deaminase | 1.65 | 1.88E-02 | Transcription regulator |
| ENSBTAG00000008250 | *SPRY4* | Protein sprouty homolog 4 | 1.64 | 2.80E-02 | Transmembrane receptor (integrin) |
| ENSBTAG00000006373 | *TSACC* | TSSK6 activating cochaperone | 0.61 | 3.89E-02 | Enzyme |
| ENSBTAG00000047238 | *ITGAM* | Integrin subunit alpha M | 1.63 | 3.06E-02 | Transporter (integrin) |
| ENSBTAG00000015549 | *PCDH18* | Protocadherin 18 | 1.62 | 4.01E-02 | Not annotated (cadherin) |
| ENSBTAG00000002885 | *MSR1* | Macrophage scavenger receptor types I and II | 1.62 | 6.52E-03 | Kinase |
| ENSBTAG00000020528 | *PCOLCE* | Procollagen C-endopeptidase enhancer | 1.61 | 2.33E-02 | Other |
| ENSBTAG00000014713 | *RARRES1* | Retinoic acid receptor responder 1 | 1.61 | 4.98E-02 | Not annotated |
| ENSBTAG00000001441 | *CARHSP1* | Calcium regulated heat stable protein 1 | 0.62 | 7.50E-03 | Other |
| ENSBTAG00000016429 | *TMEM205* | Transmembrane protein 205 | 0.62 | 3.06E-02 | Other |
| ENSBTAG00000031737 | *TMEM102* | Transmembrane protein 102 | 0.63 | 2.80E-02 | Other |
| ENSBTAG00000025778 | *EMC9* | ER membrane protein complex subunit 9 | 0.64 | 4.01E-02 | Other |
| ENSBTAG00000007071 | *RAI14* | Retinoic acid induced 14 | 1.57 | 4.01E-02 | Enzyme |

| | | | | | |
|---|---|---|---|---|---|
| ENSBTAG00000005250 | *BGN* | Biglycan | 1.57 | 3.27E-02 | Other |

| | | | | | |
|---|---|---|---|---|---|
| ENSBTAG00000038048 | *MRC1* | Mannose receptor, C type 1 | 1.57 | 4.01E-02 | Other |
| ENSBTAG00000002942 | *SLC2A10* | Solute carrier family 2 member 10 | 1.57 | 4.98E-02 | Other |
| ENSBTAG00000007156 | *AGAP2* | Arfgap with gtpase domain, ankyrin repeat and PH domain 2 | 1.55 | 7.50E-03 | Transcription regulator |
| ENSBTAG00000020144 | *GPR34* | G protein-coupled receptor 34 | 1.55 | 3.92E-02 | Other (immunoglobulin) |
| ENSBTAG00000009139 | *LDLRAD4* | Low density lipoprotein receptor class A domain containing 4 | 0.65 | 4.98E-02 | Transcription regulator |
| ENSBTAG00000021899 | *PDE9A* | Phosphodiesterase 9A | 1.54 | 1.88E-02 | Enzyme |
| ENSBTAG00000007228 | *CBX7* | Chromobox 7 | 0.65 | 3.81E-02 | Other |
| ENSBTAG00000012405 | *PEAR1* | Platelet endothelial aggregation receptor 1 | 1.54 | 3.06E-02 | Other |
| ENSBTAG00000002029 | *IGSF5* | Immunoglobulin superfamily member 5 | 0.65 | 3.28E-02 | Other (immunoglobulin) |
| ENSBTAG00000018438 | *RRAGD* | Ras-related GTP-binding protein D | 0.65 | 1.08E-02 | Transmembrane receptor (integrin) |
| ENSBTAG00000001335 | *GHR* | Growth hormone receptor | 1.53 | 3.92E-02 | Other (immunoglobulin) |
| ENSBTAG00000007187 | *INF2* | Inverted formin, FH2 and WH2 domain containing | 1.52 | 4.01E-02 | Kinase |
| ENSBTAG00000008004 | *NCF2* | Neutrophil cytosolic factor 2 | 1.52 | 4.48E-02 | Other |
| ENSBTAG00000013745 | *ITGA5* | Integrin subunit alpha 5 | 1.51 | 3.34E-02 | Enzyme |

| | | | | | |
|---|---|---|---|---|---|
| ENSBTAG00000003191 | *FSCN1* | Fascin actin-bundling protein 1 | 1.51 | 4.73E-02 | Other |
| ENSBTAG00000017664 | *HGF* | Hepatocyte growth factor | 1.51 | 4.31E-02 | Other |
| ENSBTAG00000014455 | *PDZD2* | PDZ domain containing 2 | 1.51 | 3.06E-02 | Not annotated |
| ENSBTAG00000015604 | *ZNF385A* | Zinc finger protein 385A | 1.50 | 3.06E-02 | Not annotated |
| ENSBTAG00000045834 | *RASSF7* | Ras association domain family member 7 | 0.67 | 3.28E-02 | Peptidase |
| ENSBTAG00000008389 | *HTRA1* | Serine protease HTRA1 | 1.49 | 3.89E-02 | Enzyme |
| ENSBTAG00000021746 | *ANXA5* | Annexin A5 | 1.48 | 4.26E-02 | Other |
| ENSBTAG00000031950 | *RAB3IP* (Uniprot) | RAB3A interacting protein (Rabin3) | 0.68 | 3.92E-02 | Kinase |

| | | | | | |
|---|---|---|---|---|---|
| ENSBTAG00000015478 | *SRD5A1* | 3-oxo-5-alpha-steroid 4-dehydrogenase 1 | 0.68 | 4.14E-02 | Enzyme |
| ENSBTAG00000026913 | *HHIPL1* | HHIP like 1 | 1.46 | 4.98E-02 | Transmembrane receptor |
| ENSBTAG00000005145 | *PEAK1* | Pseudopodium enriched atypical kinase 1 | 1.46 | 3.06E-02 | G-protein coupled receptor |
| ENSBTAG00000004423 | *ARHGAP42* | Rho gtpase activating protein 42 | 1.45 | 2.01E-02 | Enzyme |
| ENSBTAG00000033284 | *CHCHD7* | Coiled-coil-helix-coiled-coil-helix domain-containing protein 7 | 0.69 | 1.08E-02 | Not annotated |
| ENSBTAG00000019832 | *TGFBR2* | Transforming growth factor beta receptor 2 | 1.45 | 3.69E-02 | Other |
| ENSBTAG00000018463 | *VIM* | Vimentin | 1.45 | 4.26E-02 | Other |
| ENSBTAG00000001862 | *PPP2R2B* | Serine/threonine-protein phosphatase 2A 55 kda regulatory subunit B beta isoform | 1.45 | 2.80E-02 | Transporter |
| ENSBTAG00000005546 | *HOXB7* | Homeobox B7 | 0.69 | 3.28E-02 | Other |

| | | | | | |
|---|---|---|---|---|---|
| ENSBTAG00000008733 | *MAGED1* | MAGE family member D1 | 1.43 | 3.28E-02 | Enzyme |
| ENSBTAG00000035084 | *FZD3* (Uniprot) | Uncharacterized protein | 0.70 | 3.06E-02 | Other |
| ENSBTAG00000003629 | *ASPA* | Aspartoacylase | 1.43 | 4.98E-02 | Other |
| ENSBTAG00000015248 | *PLA2G16* (Uniprot) | Uncharacterized protein | 0.70 | 2.53E-02 | Not annotated |
| ENSBTAG00000015587 | *TTC32* | Tetratricopeptide repeat domain 32 | 0.70 | 3.92E-02 | Transmembrane receptor |
| ENSBTAG00000006777 | *TLCD1* | TLC domain containing 1 | 0.71 | 3.28E-02 | Other (cadherin) |
| ENSBTAG00000010347 | *EZR* | Ezrin | 0.71 | 3.79E-02 | Transcription regulator |
| ENSBTAG00000001956 | *HINT3* | Histidine triad nucleotide-binding protein 3 | 0.71 | 7.50E-03 | Other |
| ENSBTAG00000011954 | *SEC11C* | SEC11 homolog C, signal peptidase complex subunit | 0.71 | 2.80E-02 | Other |
| ENSBTAG00000019537 | *MARCH8* (Uniprot) | E3 ubiquitin-protein ligase MARCH8 | 1.41 | 3.06E-02 | G-protein coupled receptor |
| ENSBTAG00000017181 | *MACROD1* | O-acetyl-ADP-ribose deacetylase MACROD1 | 0.71 | 3.92E-02 | Enzyme |
| ENSBTAG00000003166 | *AXL* | AXL receptor tyrosine kinase | 1.39 | 2.80E-02 | Transporter |
| ENSBTAG00000004138 | *GLTSCR1* | Glioma tumor suppressor candidate region gene 1 | 1.39 | 3.89E-02 | Enzyme |
| ENSBTAG00000008953 | *TAP1* | Antigen peptide transporter 1 | 0.72 | 3.92E-02 | Not annotated |
| ENSBTAG00000020739 | *NXT2* | NTF2-related export protein 2 | 0.73 | 3.06E-02 | Not annotated |
| ENSBTAG00000013419 | *HTATIP2* | HIV-1 Tat interactive protein 2 | 0.74 | 3.06E-02 | Other |

| | | | | | |
|---|---|---|---|---|---|
| ENSBTAG00000044159 | *C4orf33* | Chromosome 4 open reading frame 33 | 0.74 | 3.06E-02 | Other |
| ENSBTAG00000019953 | *CYBB* | Cytochrome b-245 heavy chain | 1.35 | 3.92E-02 | Growth factor |
| ENSBTAG00000004472 | *DYNLT1* | Bos taurus dynein, light chain, Tctex-type 1 (DYNLT1), mrna. | 0.74 | 3.32E-02 | Other |
| ENSBTAG00000010351 | *SNX25* | Sorting nexin-25 | 0.75 | 7.50E-03 | Enzyme |
| ENSBTAG00000019246 | *SC5DL* (Uniprot) | Sterol-C5-desaturase (ERG3 delta-5-desaturase homolog, S. Cerevisiae)-like | 0.77 | 3.06E-02 | Other |
| ENSBTAG00000026248 | *POP7* | Ribonuclease P protein subunit p20 | 0.77 | 3.79E-02 | Enzyme |
| ENSBTAG00000021779 | *MGST2* | Microsomal glutathione S-transferase 2 | 0.77 | 4.98E-02 | Other |
| ENSBTAG00000009127 | *TSPYL4* | TSPY like 4 | 0.78 | 4.01E-02 | Other |
| ENSBTAG00000004790 | *UBE2T* | Ubiquitin-conjugating enzyme E2 T | 0.78 | 2.33E-02 | Enzyme |
| ENSBTAG00000013952 | *HNRNPD* | Heterogeneous nuclear ribonucleoprotein D | 1.28 | 4.26E-02 | G-protein coupled receptor |
| ENSBTAG00000003935 | *TMBIM1* | Transmembrane BAX inhibitor motif containing 1 | 0.79 | 1.08E-02 | Other |
| ENSBTAG00000018588 | *TMBIM6* | Bax inhibitor 1 | 0.80 | 3.06E-02 | Transporter |
| ENSBTAG00000012885 | *ACAT1* | Acetyl-coa acetyltransferase, mitochondrial | 0.81 | 3.06E-02 | Transporter |
| ENSBTAG00000001108 | *GMCL1* | Germ cell-less protein-like 1 | 0.84 | 4.33E-02 | Enzyme |

List of differentially expressed genes between pregnant and non-pregnant cows. The gene name and description are obtained with Biomart using Ensembl release 87. Where indicated the gene names were retrieved from Uniprot annotation or by homology with human genes. **FC** = Fold Change; **P-adj** = Benjamini-Hochberg adjusted P-value. IPA annotation = classification of the proteins encoded by the DE genes. Between parentesis the domain information from PFam.

***Table SM 2. GSEA enriched KEGG pathways for the genes (A) up-regulated and (B) downregulated in pregnant cows compared to non-pregnant cows.***

**A)**

| KEGG gene set | Size | NES | FDR | Core enrichment |
|---|---|---|---|---|
| ECM receptor interaction | 73 | 2.26 | ~0 | *TNC, ITGA5, COL5A2, HSPG2, COL5A3, COL4A2, SDC3, FN1, COL6A2, ITGA4, SDC2, COL6A1, THBS3, COL6A3, ITGA1, COL5A1, ITGA9, THBS2, TNXB, LAMA4, COL4A1, COL3A1, COL2A1, ITGA10, TNR, THBS1, SPP1, VWF, COL1A2, LAMC1, COL1A1, COL4A4, ITGA2B, LAMA2, SV2A, LAMB1, LAMC3,* |
| Glycosaminoglycan biosynthesis chondroitin sulfate | 18 | 1.97 | 6.97E-03 | *CHSY3, CHST3, CHST7, CHST11, XYLT1, CHST15, DSE, CHST14, CHSY1, CSGALNACT2, CHPF2, B4GALT7,* |
| Cell adhesion molecules cams | 88 | 1.91 | 1.10E-02 | *CNTN2, ITGAM, NFASC, L1CAM, SIGLEC1, CDH2, NCAM1, CNTNAP1, SDC3, NRXN2, CADM3, ITGA4, SDC2, CNTN1, ITGA9, ITGB2, NLGN1, SPN, SELPLG, CD86, CDH5, CD276, SELP, PDCD1, SELL, ESAM, ITGAL, VCAN, CD22, CD4, NCAM2, HLA-DOB, NLGN2,* |
| Focal adhesion | 174 | 1.78 | 4.10E-02 | *TNC, ITGA5, HGF, COL5A2, COL5A3, COL4A2, FN1, COL6A2, ITGA4, COL6A1, THBS3, COL6A3, ITGA1, PAK3, COL5A1, ITGA9, THBS2, FLT1, TNXB, LAMA4, COL4A1, COL3A1, KDR, COL2A1, ITGA10, TNR, PDGFRB, PARVB, PIK3CG, THBS1, SRC, TLN2, SPP1, VWF, ACTB, ZYX, COL1A2, PARVG, SHC2, PIK3R1, LAMC1, COL1A1, PXN, COL4A4, ITGA2B, SHC1, ACTN2, LAMA2, VEGFC, PRKCA, LAMB1, LAMC3, SOS1* |

| | | | | |
|---|---|---|---|---|
| Basal cell carcinoma | 35 | 1.77 | 3.49E-02 | *WNT6, WNT5A, WNT5B, WNT2, GLI3, TCF7, TP53, SMO, BMP2, DVL3, WNT7A, LEF1, WNT16, SUFU* |
| Hedgehog signaling pathway | 33 | 1.69 | 4.59E-02 | *WNT6, WNT5A, WNT5B, WNT2, GLI3, BMP6, SMO, BMP2, PRKACA, WNT7A, WNT16, SUFU* |

**Size** = number of genes in the gene set used for the analysis, **NES** = Normalized Enrichment Score, **FDR** = False Discovery Rate, **Core enrichment** =set of genes that contribute most to the enrichment result.

**B)**

| KEGG gene set | Size | NES | FDR | Core enrichment |
|---|---|---|---|---|
| Ribosome | 47 | -2.27 | ~0 | *RPL24, RPS27A, RPL15, FAU, RPS9, RPL18A, RPSA, RPL29, RPS8, RPLP0, RPS20, RPL19, RPL30, RPS2, RPS6, UBA52, RPL14, RPS28, RPS4X, RPL28, RPS3, RPL18, RPL38, RPS17, RPL7A, RPL37A, RPS7, RPL36A, RPS5, RPL34, RPS18, RPL37, RPL13, RPS16, RPL11, RPS11, RPL26, RPLP1, RPL10A, RPL36, RPS29* |
| Valine leucine and isoleucine degradation | 40 | -1.97 | 5.59E-03 | *DLD, HADH, HMGCL, ACAA2, OXCT1, ACADM, ALDH6A1, MCCC1, ACAD8, HMGCS1, PCCB, ACADS, HSD17B10, MUT, BCKDHA, IL4I1, IVD, HIBADH, HIBCH, ALDH3A2, ACAT1, EHHADH, AOX1, BCKDHB, ABAT, DBT* |
| Peroxisome | 64 | -1.86 | 1.71E-02 | *PEX19, PEX10, HMGCL, ACOT8, PEX5, PEX7, CRAT, CAT, ACSL3, MVK, ACSL4, PEX14, PEX11G, ACSL5, GSTK1, DECR2, ABCD3, HACL1, ACSL1, PRDX5, HSD17B4, ACOX3, SCP2, ACOX2, PRDX1, DDO, PEX1, EHHADH, PMVK, IDH1, CROT, PXMP2, NUDT12, SLC27A2, EPHX2* |
| Butanoate metabolism | 23 | -1.82 | 2.06E-02 | *HADH, HMGCL, OXCT1, HMGCS1, ACADS, ALDH5A1, L2HGDH, ALDH3A2, ACAT1, EHHADH, ABAT, ACSM1, ACSM3* |

**Size** = dimension of the gene set used for the analysis, **NES** = Normalized Enrichment Score, **FDR** = False Discovery Rate, **Core enrichment** = set of genes that contribute most to the enrichment result.

***Table SM 3. NET-GE GO terms enrichment results***

| GO class | GO Term | N1 | N2 | BH corrected P-value | Description |
|---|---|---|---|---|---|
| BP | GO:0022610 | 30 | 2294 | 7.40E-03 | Biological adhesion |
| BP | GO:0097485 | 19 | 1136 | 9.03E-03 | Neuron projection guidance |
| BP | GO:0090287 | 12 | 493 | 9.69E-03 | Regulation of cellular response to growth factor stimulus |
| BP | GO:0051240 | 25 | 1916 | 1.10E-02 | Positive regulation of multicellular organismal process |
| BP | GO:0007411 | 19 | 1136 | 1.13E-02 | Axon guidance |
| BP | GO:0042476 | 9 | 294 | 1.17E-02 | Odontogenesis |
| BP | GO:0017015 | 8 | 241 | 1.20E-02 | Regulation of transforming growth factor beta receptor signaling pathway |
| BP | GO:1902533 | 26 | 2106 | 1.23E-02 | Positive regulation of intracellular signal transduction |
| BP | GO:0043062 | 17 | 991 | 1.23E-02 | Extracellular structure organization |
| BP | GO:0006897 | 17 | 1025 | 1.25E-02 | Endocytosis |
| BP | GO:1901568 | 8 | 240 | 1.25E-02 | Fatty acid derivative metabolic process |
| BP | GO:0030512 | 7 | 165 | 1.26E-02 | Negative regulation of transforming growth factor beta receptor signaling pathway |
| BP | GO:0098602 | 15 | 846 | 1.29E-02 | Single organism cell adhesion |
| BP | GO:0030198 | 17 | 986 | 1.32E-02 | Extracellular matrix organization |
| BP | GO:0001503 | 11 | 397 | 1.33E-02 | Ossification |
| BP | GO:0006690 | 8 | 240 | 1.35E-02 | Icosanoid metabolic process |
| BP | GO:0007155 | 30 | 2287 | 1.39E-02 | Cell adhesion |
| BP | GO:0031589 | 10 | 410 | 1.62E-02 | Cell-substrate adhesion |
| BP | GO:0031100 | 6 | 136 | 1.74E-02 | Organ regeneration |
| BP | GO:0033559 | 8 | 268 | 1.89E-02 | Unsaturated fatty acid metabolic process |

| | | | | | |
|---|---|---|---|---|---|
| BP | GO:0042475 | 7 | 211 | 2.30E-02 | Odontogenesis of dentin-containing tooth |
| BP | GO:0032570 | 5 | 94 | 2.39E-02 | Response to progesterone |
| BP | GO:0007159 | 5 | 96 | 2.41E-02 | Leukocyte cell-cell adhesion |
| BP | GO:0030574 | 6 | 152 | 2.43E-02 | Collagen catabolic process |
| BP | GO:0046098 | 2 | 4 | 2.47E-02 | Guanine metabolic process |
| BP | GO:0048660 | 7 | 223 | 2.64E-02 | Regulation of smooth muscle cell proliferation |
| BP | GO:0045597 | 25 | 2141 | 2.68E-02 | Positive regulation of cell differentiation |
| BP | GO:0044236 | 7 | 221 | 2.70E-02 | Multicellular organismal metabolic process |
| BP | GO:0060394 | 3 | 23 | 2.71E-02 | Negative regulation of pathway-restricted SMAD protein phosphorylation |
| BP | GO:0070661 | 7 | 227 | 2.75E-02 | Leukocyte proliferation |
| BP | GO:0001952 | 7 | 229 | 2.81E-02 | Regulation of cell-matrix adhesion |
| BP | GO:0009967 | 32 | 3132 | 2.83E-02 | Positive regulation of signal transduction |
| BP | GO:0007568 | 12 | 659 | 2.84E-02 | Aging |
| BP | GO:0032963 | 6 | 176 | 3.03E-02 | Collagen metabolic process |
| BP | GO:0006898 | 10 | 495 | 3.06E-02 | Receptor-mediated endocytosis |
| BP | GO:0031099 | 8 | 322 | 3.10E-02 | Regeneration |
| BP | GO:0006145 | 2 | 6 | 3.14E-02 | Purine nucleobase catabolic process |
| BP | GO:0048661 | 5 | 117 | 3.14E-02 | Positive regulation of smooth muscle cell proliferation |
| BP | GO:0044243 | 6 | 178 | 3.15E-02 | Multicellular organismal catabolic process |
| BP | GO:0071902 | 13 | 799 | 3.16E-02 | Positive regulation of protein serine/threonine kinase activity |
| BP | GO:0030334 | 21 | 1708 | 3.21E-02 | Regulation of cell migration |
| BP | GO:0007567 | 3 | 27 | 3.27E-02 | Parturition |
| BP | GO:0050730 | 11 | 583 | 3.28E-02 | Regulation of peptidyl-tyrosine phosphorylation |
| BP | GO:0048646 | 27 | 2487 | 3.31E-02 | Anatomical structure formation involved in morphogenesis |
| BP | GO:0043406 | 11 | 583 | 3.37E-02 | Positive regulation of MAP kinase activity |

| | | | | | |
|---|---|---|---|---|---|
| BP | GO:0016337 | 13 | 778 | 3.40E-02 | Single organismal cell-cell adhesion |
| BP | GO:0008283 | 25 | 2213 | 3.44E-02 | Cell proliferation |
| BP | GO:0051347 | 21 | 1754 | 3.72E-02 | Positive regulation of transferase activity |
| BP | GO:0001837 | 5 | 123 | 3.78E-02 | Epithelial to mesenchymal transition |
| BP | GO:0009100 | 6 | 189 | 3.88E-02 | Glycoprotein metabolic process |
| BP | GO:0031401 | 27 | 2590 | 3.97E-02 | Positive regulation of protein modification process |
| BP | GO:0090288 | 7 | 271 | 4.01E-02 | Negative regulation of cellular response to growth factor stimulus |
| BP | GO:0008366 | 6 | 199 | 4.03E-02 | Axon ensheathment |
| BP | GO:2000145 | 21 | 1795 | 4.04E-02 | Regulation of cell motility |
| BP | GO:0044259 | 6 | 196 | 4.04E-02 | Multicellular organismal macromolecule metabolic process |
| BP | GO:0010811 | 7 | 266 | 4.08E-02 | Positive regulation of cell-substrate adhesion |
| BP | GO:0007272 | 6 | 199 | 4.09E-02 | Ensheathment of neurons |
| BP | GO:0044246 | 5 | 130 | 4.09E-02 | Regulation of multicellular organismal metabolic process |
| BP | GO:0001525 | 13 | 829 | 4.11E-02 | Angiogenesis |
| BP | GO:0007229 | 6 | 195 | 4.15E-02 | Integrin-mediated signaling pathway |
| BP | GO:0033674 | 19 | 1531 | 4.16E-02 | Positive regulation of kinase activity |
| BP | GO:0043627 | 10 | 533 | 4.16E-02 | Response to estrogen |
| BP | GO:0044272 | 7 | 268 | 4.19E-02 | Sulfur compound biosynthetic process |
| BP | GO:0042327 | 25 | 2349 | 4.49E-02 | Positive regulation of phosphorylation |
| BP | GO:0001934 | 23 | 2079 | 4.52E-02 | Positive regulation of protein phosphorylation |
| BP | GO:0006691 | 4 | 80 | 4.54E-02 | Leukotriene metabolic process |
| BP | GO:0060393 | 4 | 82 | 4.56E-02 | Regulation of pathway-restricted SMAD protein phosphorylation |
| BP | GO:0002688 | 6 | 209 | 4.60E-02 | Regulation of leukocyte chemotaxis |
| BP | GO:0010810 | 9 | 458 | 4.62E-02 | Regulation of cell-substrate adhesion |
| BP | GO:0050878 | 20 | 1711 | 4.63E-02 | Regulation of body fluid levels |

| | | | | | |
|---|---|---|---|---|---|
| BP | GO:0006954 | 16 | 1209 | 4.63E-02 | Inflammatory response |
| BP | GO:0043410 | 15 | 1094 | 4.64E-02 | Positive regulation of MAPK cascade |
| BP | GO:0019369 | 5 | 140 | 4.65E-02 | Arachidonic acid metabolic process |
| BP | GO:0045860 | 18 | 1468 | 4.87E-02 | Positive regulation of protein kinase activity |
| BP | GO:0000187 | 8 | 378 | 4.91E-02 | Activation of MAPK activity |
| MF | GO:0004464 | 3 | 6 | 1.64E-03 | Leukotriene-C4 synthase activity |
| MF | GO:0005539 | 12 | 607 | 2.06E-02 | Glycosaminoglycan binding |
| MF | GO:0034713 | 3 | 16 | 2.21E-02 | Type I transforming growth factor beta receptor binding |
| MF | GO:1901681 | 12 | 664 | 3.54E-02 | Sulfur compound binding |
| MF | GO:0019239 | 4 | 62 | 3.86E-02 | Deaminase activity |
| CC | GO:0009986 | 26 | 1474 | 2.02E-04 | Cell surface |
| CC | GO:0044459 | 48 | 5012 | 1.66E-02 | Plasma membrane part |
| CC | GO:0005887 | 33 | 2884 | 1.85E-02 | Integral component of plasma membrane |
| CC | GO:0044420 | 9 | 340 | 1.94E-02 | Extracellular matrix part |
| CC | GO:0031012 | 14 | 799 | 1.97E-02 | Extracellular matrix |
| CC | GO:0005578 | 13 | 639 | 2.31E-02 | Proteinaceous extracellular matrix |
| CC | GO:0070161 | 16 | 1072 | 3.70E-02 | Anchoring junction |

**GO Class** = GO term category: Biological Process (BP), Cellular Component (CC) or Molecular Function (MF); **N1** = **N1 -** The number of input genes associated to the GO term, **N2** = The number of genes associated to the GO term; **BH P-value** = Benjamini-Hochberg adjusted P-value.

***Table SM 4. Functional classification of the genes by IPA®***

| Biological category | BH P-value | Molecules |
|---|---|---|
| Cellular Movement | 6.3E-05-2.88E-02 | HTATIP2, MSR1, FZD3, RARRES1, SCUBE3, GPR34, TMBIM6, LOXL2, SPRY4, TGFBR2, TMEM102, ANXA5, HGF, EZR, CYBB, POSTN, PLA2G16, PEAK1, TGFBI, BGN, VIM, ITGA5, AGAP2, ITGAM, LDLRAD4, FSCN1, NCF2, ACAT1, PTGS2, ALOX5, HTRA1 |

| | | |
|---|---|---|
| Cellular Function and Maintenance | 1.45E-04-2.88E-02 | MRC1, PEAK1, MSR1, VIM, TGFBR2, GHR, ITGAM, HGF, ANXA5, EZR, CYBB, PEAR1, GRIK2, PTGS2 |
| Free Radical Scavenging | 3.99E-04-2.88E-02 | MRC1, ITGAM, MSR1, HGF, NCF2, CYBB, ITGA5, TMBIM6, PTGS2, ALOX5 |
| Small Molecule Biochemistry | 3.99E-04-2.88E-02 | MRC1, PLA2G16, HTATIP2, MSR1, QRFPR, SRD5A1, CYP4F3, ITGA5, VIM, ASPA, TGFBR2, TGM1, ITGAM, GHR, TMEM102, MGST2, ANXA5, HGF, NCF2, CYBB, ACAT1, PTGS2, ALOX5 |
| Cell Morphology | 3.99E-04-2.88E-02 | ZNF385A, GDA, MSR1, COL8A1, GPR34, TMBIM6, DYNLT1, TAP1, LOXL2, PCOLCE, SPRY4, TGFBR2, HGF, EZR, ANXA5, CYBB, POSTN, PEAK1, ADD2, TGFBI, PDE9A, BGN, ITGA5, VIM, MANBA, AGAP2, ITGAM, GHR, NPR3, FSCN1, GMCL1, PTGS2, HTRA1 |
| Cell-To-Cell Signaling and Interaction | 1.78E-03-2.88E-02 | MRC1, PEAK1, ADD2, MSR1, TGFBI, BGN, VIM, ITGA5, TAP1, TGFBR2, GHR, ITGAM, EZR, HGF, ANXA5, CYBB, POSTN, PEAR1, GRIK2, PTGS2 |
| Cellular Development | 2.53E-03-2.88E-02 | CBX7, MAGED1, ZNF385A, BGN, ITGA5, VIM, SCUBE3, AGAP2, LOXL2, GPC5, TGFBR2, TGM1, NPR3, HOXB7, EZR, FSCN1, HGF, POSTN, PTGS2, ALOX5, HTRA1 |
| Cellular Growth and Proliferation | 2.53E-03-2.88E-02 | CBX7, MAGED1, BGN, COL8A1, VIM, ITGA5, SCUBE3, AGAP2, GPC5, TGFBR2, GHR, TGM1, NPR3, HOXB7, FSCN1, EZR, HGF, POSTN, PTGS2, ALOX5, HTRA1 |
| Cell Death and Survival | 4.04E-03-2.88E-02 | CBX7, HTATIP2, MAGED1, ZNF385A, PCDH18, GDA, MSR1, FZD3, WNT6, TMBIM6, TAP1, LOXL2, TGFBR2, MARCH8, HGF, EZR, ANXA5, POSTN, CYBB, GRIK2, RAI14, PLA2G16, ADD2, TGFBI, PDE9A, BGN, VIM, ITGA5, AGAP2, ASPA, KCNB1, TGM1, ITGAM, GHR, PPP2R2B, NCF2, GMCL1, ACAT1, PTGS2, ALOX5, HTRA1 |
| Cellular Compromise | 5.05E-03-2.88E-02 | TGFBR2, ITGAM, TGM1, ADD2, ANXA5, CYBB, VIM, ASPA |
| Lipid Metabolism | 5.05E-03-2.88E-02 | MRC1, PLA2G16, MSR1, QRFPR, SRD5A1, CYP4F3, VIM, ASPA, TGFBR2, GHR, MGST2, ANXA5, HGF, ACAT1, PTGS2, ALOX5 |
| Cellular Assembly and Organization | 7.66E-03-2.88E-02 | SCG3, PEAK1, MSR1, ADD2, BGN, VIM, TMBIM6, PCOLCE, GHR, EZR, HGF, ANXA5, FSCN1, PTGS2 |
| Cell Cycle | 8.7E-03-2.88E-02 | TGFBR2, CBX7, GHR, HGF |

| | | |
|---|---|---|
| Molecular Transport | 8.79E-03-2.88E-02 | MRC1, TGFBR2, GHR, MSR1, SRD5A1, HGF, NCF2, ACAT1, CYBB, PTGS2, ALOX5 |
| Protein Synthesis | 1.26E-02-2.88E-02 | MRC1, GHR, NPR3, HGF, PTGS2, TMBIM1 |
| Carbohydrate Metabolism | 1.93E-02-2.88E-02 | HTATIP2, ANXA5 |
| Vitamin and Mineral Metabolism | 2.88E-02-2.88E-02 | GHR, QRFPR, SRD5A1, BGN, HGF, ACAT1, VIM, ASPA |
| Amino Acid Metabolism | 2.88E-02-2.88E-02 | TGFBR2, GHR, TGM1, TMEM102, HGF, ITGA5 |
| Post-Translational Modification | 2.88E-02-2.88E-02 | TGFBR2, GHR, TGM1, TMEM102, HGF, ITGA5 |
| Drug Metabolism | 2.88E-02-2.88E-02 | SRD5A1, HGF, PTGS2 |
| DNA Replication, Recombination, and Repair | 2.88E-02-2.88E-02 | VIM |
| Protein Trafficking | 2.88E-02-2.88E-02 | HGF |

**BH P-value**: (P-values after multiple testing correction by using Benjamini–Hochberg method).

***Table SM 5. Upstream regulator analysis by IPA®***

| Upstream Regulator | Fold Change | Molecule Type | P-value of overlap | Target molecules in dataset |
|---|---|---|---|---|
| progesterone | | chemical - endogenous mammalian | 1.08E-07 | ACAT1, EZR, GHR, HGF, ITGA5, LDLRAD4, MRC1, POSTN, PTGS2, SRD5A1, STEAP1, TGFBI, TGM1, VIM, WNT6 |
| IGF1 | 0.34 | growth factor | 4.03E-07 | BGN, CYBB, GHR, HGF, ITGA5, NCF2, PCOLCE, PTGS2, SRD5A1, TAP1, TGFBI, VIM |
| SP1 | 0.02 | transcription regulator | 1.03E-05 | ALOX5, COL8A1, EZR, GHR, HGF, ITGAM, MRC1, NCF2, PTGS2, TGFBR2, TGM1, VIM |
| APOE | 0.46 | transporter | 1.28E-05 | ACAT1, BGN, CYBB, ITGA5, ITGAM, MSR1, NCF2, PTGS2 |
| IL10RA | 0.20 | transmembrane receptor | 1.48E-05 | ALOX5, CLEC12A, GDA, GHR, HGF, PCOLCE, PLA2G16, POSTN, TAP1 |
| HIF1A | -0.09 | transcription regulator | 1.60E-05 | BGN, CYP4F3, EMC9, FSCN1, GHR, ITGA5, LOXL2, PTGS2, SRD5A1, VIM |
| HNRNPAB | -0.04 | enzyme | 2.11E-05 | HTRA1, PTGS2, VIM |
| CBP-ICSBP-IRF-1-PU.1 | | complex | 2.16E-05 | CYBB, NCF2 |
| tretinoin | | chemical - endogenou | 2.39E-05 | ALOX5, ANXA5, CYBB, CYP4F3, GHR, GRIK2, HOXB7, ITGAM, LOXL2, NCF2, |

| | | | | |
|---|---|---|---|---|
| | | s mammalian | | POSTN, PTGS2, RAI14, RARRES1, TAP1, TGFBI, TGFBR2, TGM1, VIM, WNT6 |
| EDN1 | -0.11 | cytokine | 4.01E-05 | ANXA5, CYBB, EZR, ITGA5, ITGAM, PTGS2, VIM |
| AGT | | growth factor | 4.58E-05 | BGN, CYBB, HGF, ITGA5, MSR1, NCF2, NPR3, POSTN, PTGS2, TGFBR2 |
| FOS | -0.09 | transcription regulator | 4.62E-05 | ADD2, EZR, FZD3, GRIK2, HTATIP2, NCF2, PCDH19, PTGS2, TGFBI, VIM, WNT6 |
| P38 MAPK | | group | 4.79E-05 | ANXA5, BGN, CYBB, CYP4F3, HTRA1, ITGAM, KCNB1, POSTN, PTGS2 |
| OLR1 | -0.12 | transmembrane receptor | 6.27E-05 | CYBB, ITGAM, NCF2, TGFBR2 |
| SP3 | 0.026 | transcription regulator | 6.75E-05 | GHR, HGF, NCF2, PTGS2, TGFBR2, TGM1, VIM |
| SPI1 | 0.559 | transcription regulator | 7.15E-05 | CYBB, ITGA5, ITGAM, MRC1, NCF2, PTGS2, VIM |
| PTGES | -0.352 | enzyme | 1.10E-04 | CYBB, EZR, PTGS2, VIM |
| BIRC5 | 0.139 | other | 1.27E-04 | BGN, COL8A1, ITGA5, VIM |
| Tgf beta | | group | 1.31E-04 | BGN, FSCN1, ITGA5, POSTN, PTGS2, TGFBI, VIM |
| PDGF BB | | complex | 1.31E-04 | GDA, GHR, ITGA5, MGST2, NPR3, PCOLCE, POSTN, PTGS2 |
| FABP4 | | transporter | 1.42E-04 | ACAT1, MSR1, PTGS2 |
| AMPK | | complex | 2.31E-04 | CYBB, FSCN1, KCNB1, PTGS2, VIM |
| Alpha catenin | | group | 2.40E-04 | BGN, ITGA5, ITGAM, PTGS2, VIM |
| IL27 | | cytokine | 3.49E-04 | ITGAM, MRC1, PTGS2, TAP1, VIM |
| TNF | -0.279 | cytokine | 4.03E-04 | ALOX5, BGN, CHSY3, CYBB, FSCN1, GHR, HGF, ITGA5, ITGAM, MGST2, MSR1, NCF2, PLA2G16, POSTN, PTGS2, RASSF7, TAP1, TGFBR2, VIM |

Top 25 upstream regulators obtained for the differential expression profile between pregnant and non-pregnant cows.

***Table SM 6 Composition of media used for sperm holding- and washing (Sperm TALP), in-vitro maturation (IVM), in-vitro fertilization (IVF) and in-vitro embryo culture (IVC)***

| **Ingredients** | **Sperm TALP** | **IVM** | **IVF** | **IVC** | **Sigma cat. no.** |
|---|---|---|---|---|---|
| **NaCl** | 5.8 g/L | | 6.7 g/L | 6.3 g/L | S5886 |
| **KCl** | 0.2 g/L | | 0.2 g/L | 0.5 g/L | S5405 |
| **NaH2PO4** | 0.04 g/L | | 0.04 g/L | | S5011 |
| **CaCl2, 2H2O** | 0.3 g/L | | 0.3 g/L | 0.3 g/L | C7902 |
| **MgCl2** | 0.2 g/L | | 0.1 g/L | | M2393 |
| **Lactic acid** | 3.7 mL/L | | 1.9 mL/L | | L4263 |

| | | | | | |
|---|---|---|---|---|---|
| **Hepes** | 1.4 g/L | | | | H3784 |
| **Hepes-acid** | 1.1 g/L | | | | H4034 |
| **NaHCO3** | 2.1 g/L | | 2.1 g/L | 2.1 g/L | S4019 |
| **Pyruvic acid** | 110 mg/L | | 0.02 g/L | | P3662 |
| **Gentamycin** | 100 mg/L | 0.04 mg/mL | | 0.05 g/L | G1264 |
| **Heparin** | | | 5 IU/mL | | H3149 |
| **Sodium disulfite** | | | 0.04 µg/mL | | S9000 |
| **Penicillamine** | | | 4.4 µg/mL | | P4875 |
| **Hypotaurine** | | | 0.16 µg/mL | | H1384 |
| **Epinephrine** | | | 0.27 µg/mL | | E4250 |
| **KH2PO4** | | | | 0.2 g/L | P5655 |
| **MgSO4** | | | | 0.2 g/L | M2643 |
| **Sodium lactate solution** | | | | 0.6 mL/mL | L4263 |
| **Sodium pyruvate** | | | | 0.08 g/L | P3662 |
| **Citric acid** | | | | 0.1 g/L | C3434 |
| **Myo-inositol** | | | | 0.5 g/L | I7508 |
| **BME amino acids** | | | | 30 µL /mL | B6766 |
| **MEM amino acids** | | | | 10 µL /mL | M7145 |
| **L-glutamine** | | 0.10 mg/mL | | 0.03 g/L | G8540 |
| **PMSG** | | 8.7 IU/mL | | | |
| **hCG** | | 4.3 IU/mL | | | |
| **Cattle Serum** | | 13% | | 5% | |
| **Dissolved in Medium 199** | | X | | | M2154 |
| **Dissolved in H2O** | X | | X | X | W1503 |
| **Osmolarity** | 280 ± 8 mOsm | 280 ± 8 mOsm | 280 ± 8 mOsm | 280 ± 8 mOsm | |

BME: Basal Medium Eagle, MEM: Minimum Essential Medium, PMSG: Pregnant Mare Serum Gonadotropin, hCG: Human Chorionic Gonadotropin

### Supplementary Figures

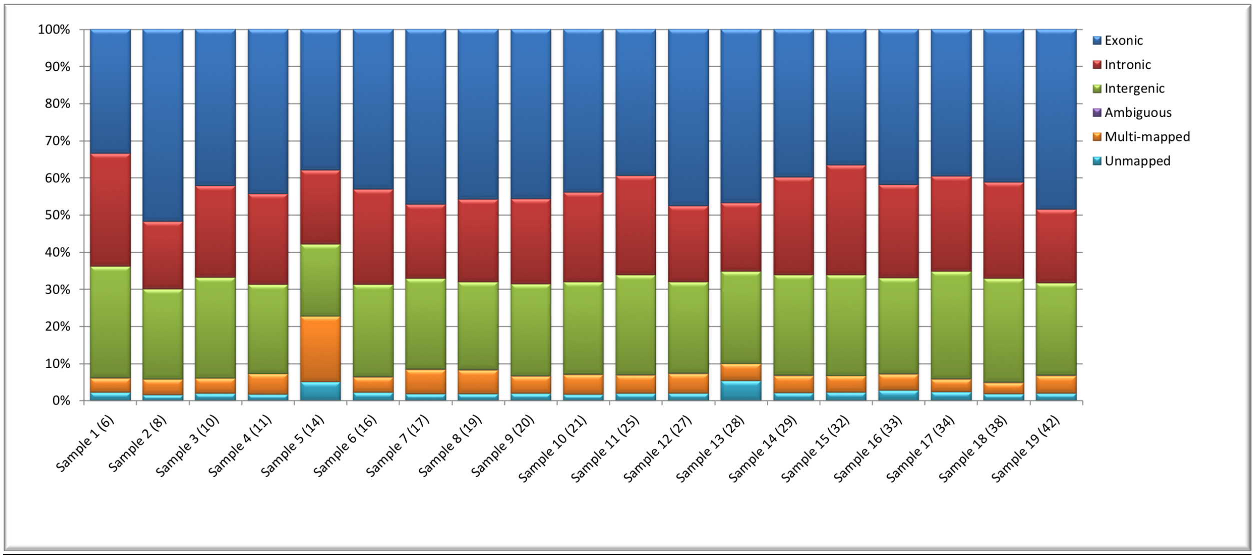


***Fig. SM 1: Quality Control after alignment***

Percentages of reads unmapped, mapped to multiple position and uniquely mapped reads classified as ambiguous exonic intronic and intergenic for each sample. Numbers in parentheses correspond to the animals' IDs.

E1: GEN161123_K 6_P01_F

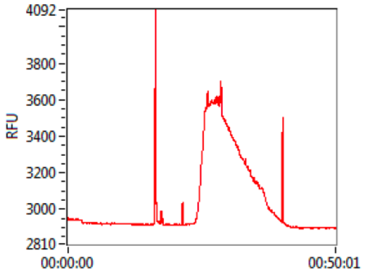


F1: GEN161123_K 7_P01_F

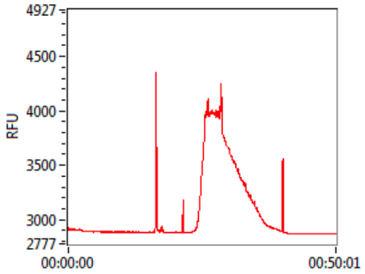


G1: GEN161123_K 8_P01_F

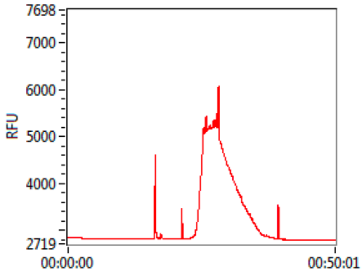


H1: GEN161123_K 10_P01_F

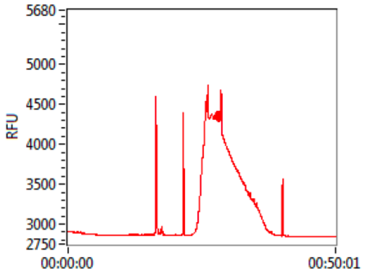


E2: GEN161123_K 11_P01_F

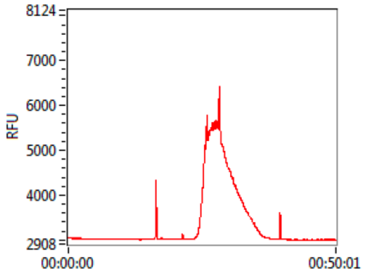


F2: GEN161123_K 12_P01_F

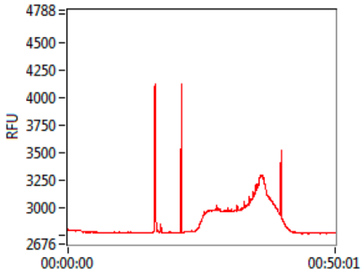


G2: GEN161123_K 14_P01_F

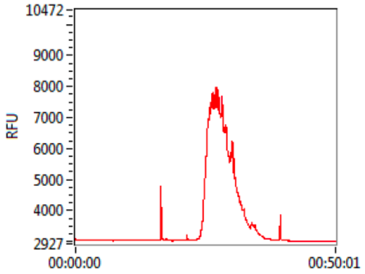


H2: GEN161123_K 16_P01_F

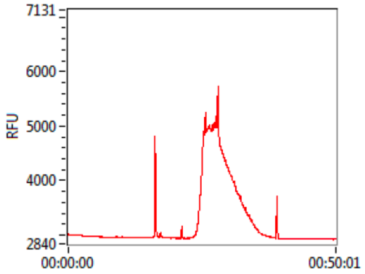


E3: GEN161123_K 17_P01_F

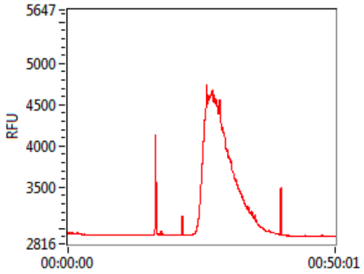


F3: GEN161123_K 18_P01_F

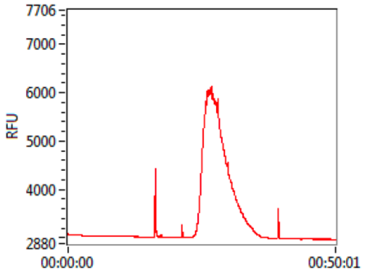


G3: GEN161123_K 19_P01_F

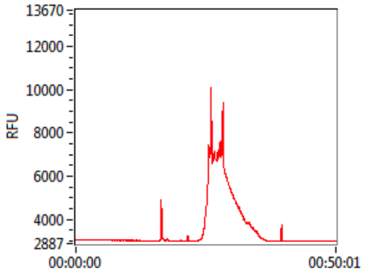


H3: GEN161123_K 20_P01_F

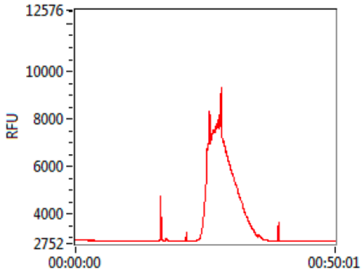

E4: GEN161123_K 21_P01_F

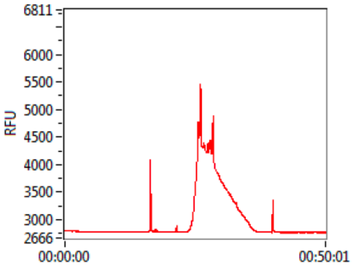


F4: GEN161123_K 25_P01_F

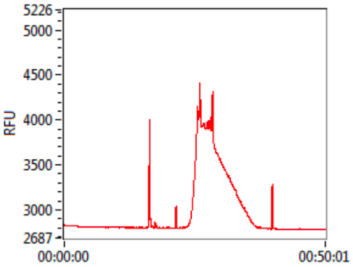


G4: GEN161123_K 27_P01_F

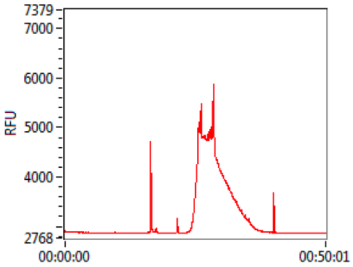


H4: GEN161123_K 28_P01_F

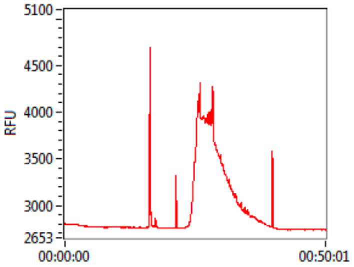


E5: GEN161123_K 29_P01_F

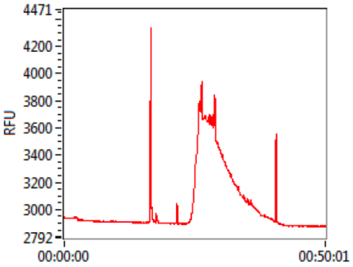


F5: GEN161123_K 32_P01_F

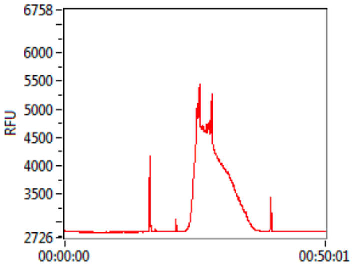


G5: GEN161123_K 33_P01_F

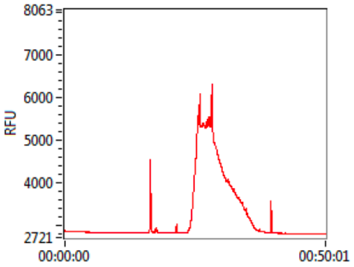


H5: GEN161123_K 34_P01_F

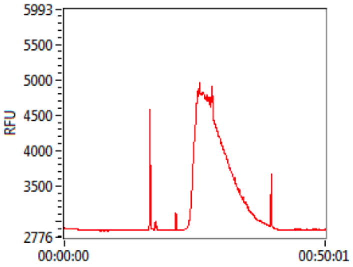


E6: GEN161123_K 38_P01_F

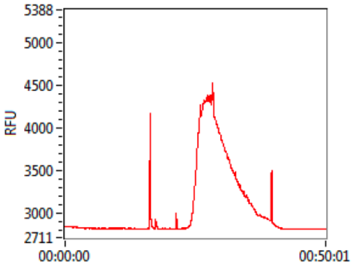


F6: GEN161123_K 42_P01_F

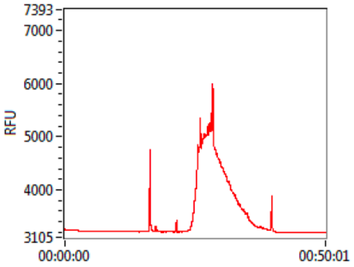


G6: GEN161123_K 43_P01_F

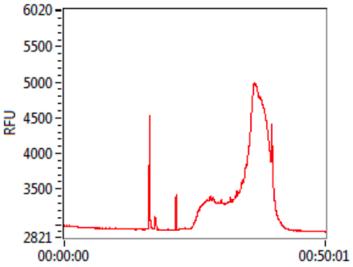

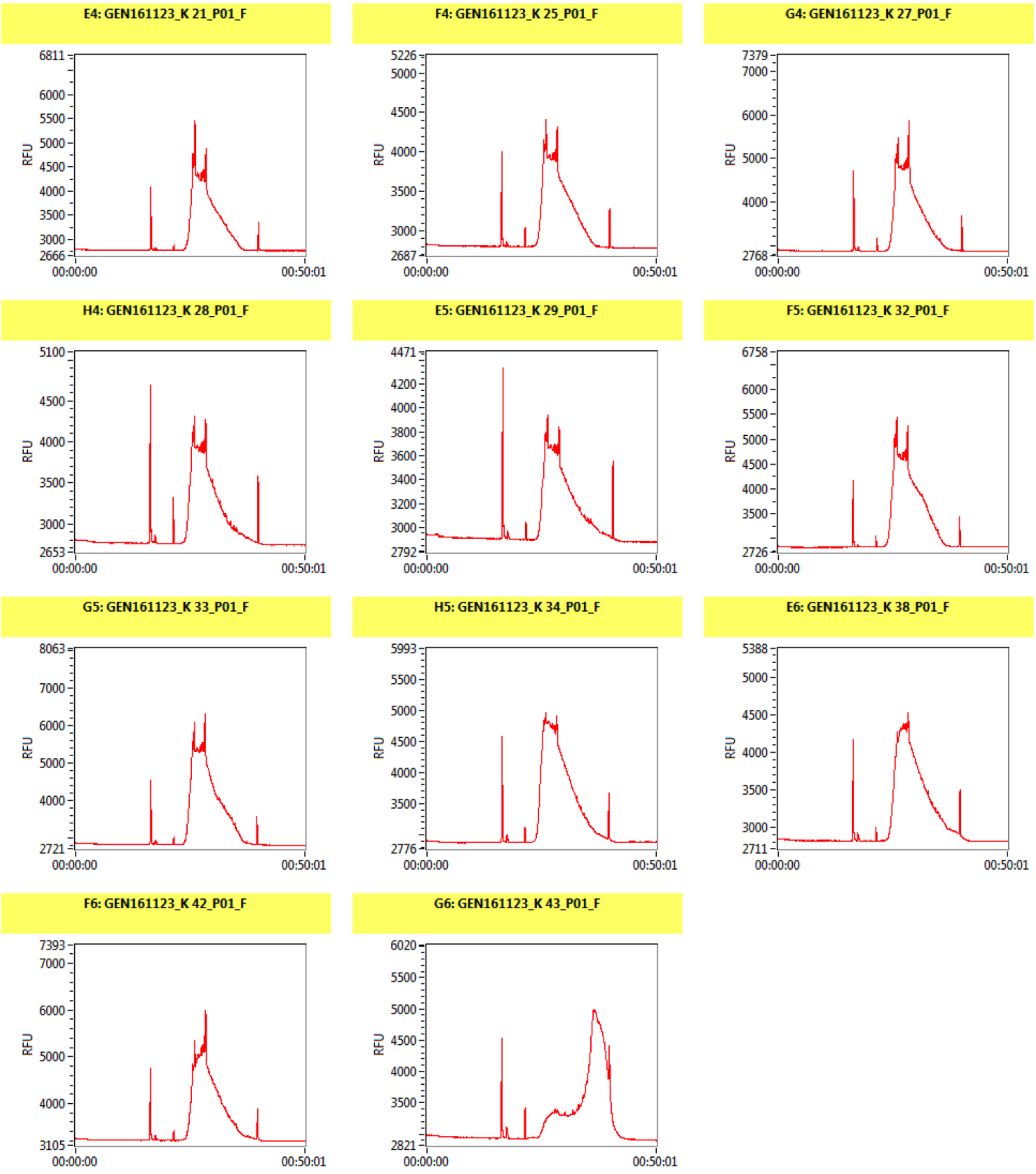


***Fig. SM 2: Library fragmentation profile***

Library fragmentation profiles. The samples G6 and F2 show a different fragmentation profile.

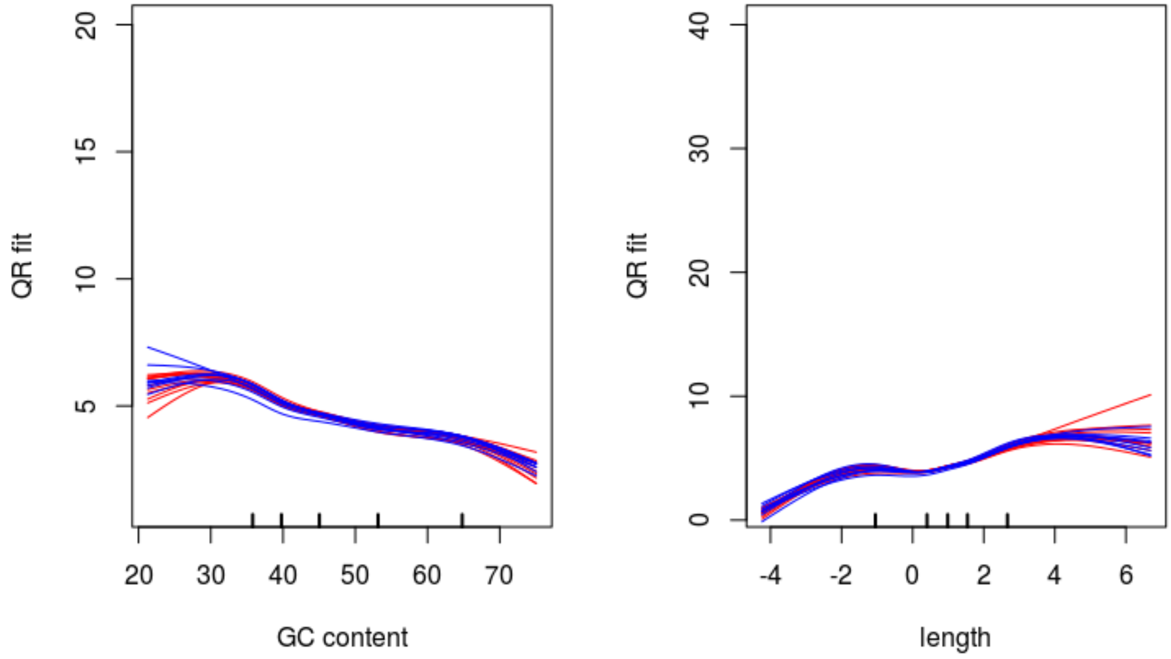


***Fig. SM 3: GC content and length bias effect across samples***

Plots of the estimated systematic GC bias and length bias effect (cqn R package) across samples. To be noted there is no relevant difference of both the GC and the length bias effect among samples.

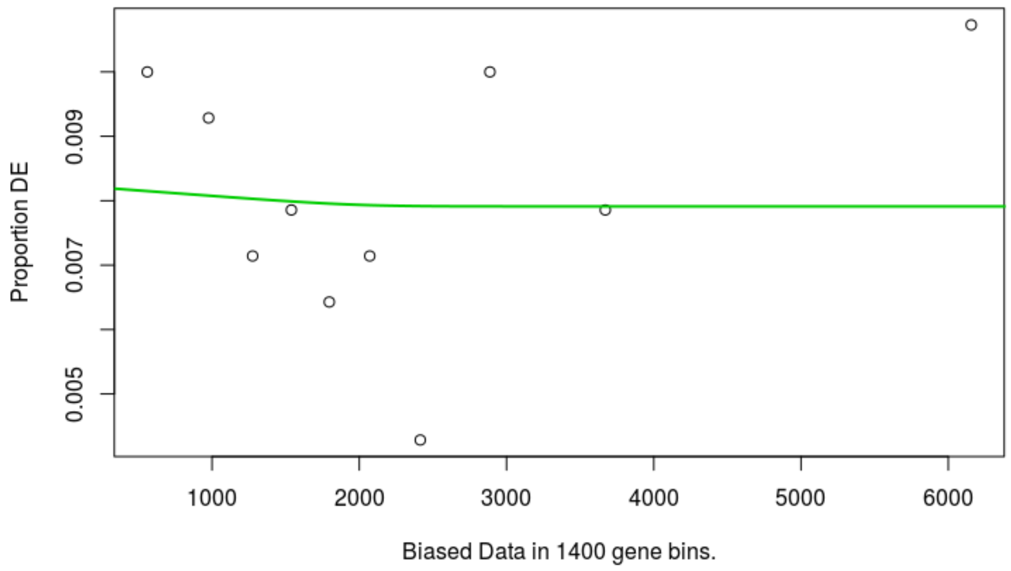


***Fig. SM 4: Length bias effect on the proportion of DE genes***

Proportion of DE genes (in a gene bin) plotted as a function of the gene length (average length of the genes in the bin). To be noted, in our results there is no relevant length bias effect on the proportion of DE genes.
